# Development of an Intranasally- and Intramuscularly-Administrable Replicon Vaccine Efficacious Against H5N1 Influenza Virus

**DOI:** 10.1101/2025.03.31.646478

**Authors:** Wynton D. McClary, Devin S. Brandt, Madeleine F. Jennewein, Jasneet Singh, Samuel Beaver, Matthew R. Ykema, Christopher Press, Eduard Melief, Julie Bakken, Pauline Fusco, Ethan Lo, Peter Battisti, Noah Cross, Darshan N. Kasal, Airn Tolnay Hartwig, Corey Casper, Richard A. Bowen, Alana Gerhardt, Emily A. Voigt

## Abstract

The risk of a respiratory viral pandemic is significant, including from the now widespread panzootic H5N1 influenza virus, highlighting the need for effective, stable, and inexpensive vaccine technologies that elicit strongly protective immunity. Intranasal vaccines can stimulate local immune responses at the site of natural respiratory viral infection, a key characteristic that can not only reduce morbidity and mortality caused by respiratory viruses but also potentially reduce viral transmissibility to limit outbreaks. Nucleic acid vaccines are now a valuable tool in pandemic responses, with high potency and rapid adaptability to target circulating or emerging viral strains; however, data are limited on which vaccine attributes are needed for efficient transmucosal delivery and immune stimulation following intranasal delivery. To demonstrate proof of concept, here we have developed a replicon vaccine expressing an H5 influenza antigen that uses a nanostructured lipid carrier (NLC) delivery system. A relationship was established between the molar ratio of positive charges on the NLC to the negative charges on the nucleic acid (N:P ratio) and the immunogenicity of the vaccine formulations, with higher N:P ratios resulting in an increase in vaccine immunogenicity. We demonstrated the ability of this replicon vaccine to be administered via intramuscular and intranasal routes with a singular vaccine formulation. The vaccine induced systemic immunity when dosed intramuscularly or intranasally in an immunocompetent mouse model, whereas intranasal dosing uniquely stimulated a strong mucosal immune response. Moreover, a mixed intramuscular/intranasal dosing strategy using this unified formulation stimulated a balanced systemic and mucosal immune response. Finally, we demonstrated the protective efficacy of this intranasally and intramuscularly/intranasally delivered H5 replicon-NLC vaccine against morbidity and mortality in a lethal H5N1 influenza challenge ferret model. This work establishes the replicon-NLC vaccine platform as a potential novel intranasal technology for rapid pandemic response.

## INTRODUCTION

Respiratory viruses pose a critical and ongoing threat to global health and economic stability [1]. An example of this is the ever-present pandemic risk posed by panzootic strains of influenza virus, such as the H5N1 strains prevalent in domestic animal populations that are driving increased infection events in humans [2–5]. The ease of transmission of respiratory viruses, coupled with their ability to mutate rapidly, makes it difficult to develop effective vaccines that keep up with this constantly evolving threat [6,7]. Furthermore, most currently approved vaccines against respiratory viruses are given via intramuscular (IM) administration. While these vaccines induce strong systemic immunity, a vaccine against a respiratory virus that is administered intranasally (IN) to establish immunity in the respiratory mucosa may not only provide protection from significant morbidity but also aid in limiting viral transmission and have the added benefit of potentially increasing vaccine uptake among the population [8–10]. To respond to the ever-present threat of the next respiratory pandemic, novel vaccine technologies are required that (1) can be developed and deployed rapidly, (2) can elicit potent immune responses at the site of infection to decrease transmissibility of the virus, and (3) are in a presentation conducive to increased uptake and patient compliance.

As was evidenced by their use during the SARS-CoV-2 global pandemic, vaccines containing antigen-expressing RNA have proven to be an especially effective tool for pandemic response due to their rapid adaptability [11,12]. However, these current RNA vaccine technologies typically involve complex manufacturing processes required to encapsulate mRNA within lipid nanoparticles (LNPs), increasing difficulty of technology transfer to global manufacturers [13,14]. Additionally, and of key importance, these RNA vaccines have not demonstrated the ability to prevent mild respiratory infections and interrupt viral transmission by vaccinated individuals [15–17].

The prevention of disease transmission may require induction of mucosal immunity at the site of natural infection (i.e., the nasal mucosa) [18,19]. This is most readily achievable through the IN administration of a vaccine. Direct delivery of a vaccine antigen to mucosal tissues increases antigen-specific mucosal IgA levels at the natural site of respiratory viral infection, which is fundamental to an effective first-line immune response [20–22]. While mucosal immunity may be key to prevent infection and spread, systemic immune responses are also critical to attenuate disease symptoms and progression [23]. Intranasally administered vaccines must therefore be formulated to not only be well tolerated when delivered to sensitive mucosal tissues but also to ensure a balanced systemic and mucosal response. Heterologous IM/IN dosing regimens may also represent an effective way to achieve this balance [24,25]. However, no nucleic acid vaccine yet exists that allows for both IM and IN routes of administration with a single formulation.

We previously developed a vaccine platform that utilizes a replicon in combination with a nanostructured lipid carrier (NLC) as a delivery vector [26–28]. NLCs can be manufactured independently of the replicon vaccine component, and the final replicon-NLC vaccine drug product is generated via simple one-step mixing of the replicon and NLC bulk drug substances.

This results in the formation of a protective complex wherein the replicon nucleic acid is electrostatically bound to the NLC surface. Further, this enables independent stockpiling of the individual vaccine components (NLC and replicon), avoiding undesirable nucleic acid degradation and allowing for greater overall flexibility for rapid vaccine manufacturing. In contrast, commonly used lipid nanoparticle (LNP)-based RNA delivery vectors require the direct incorporation of RNA into the LNP reaction mixture in order to fully encapsulate the RNA within the LNP core, exposing the RNA to harsh processing during LNP manufacture (e.g., elevated temperatures and high-shear mixing) [13]. Furthermore, the NLC composition itself does not need to be fine-tuned to the physical properties (i.e., the size) of each replicon construct to be effective, further enhancing the flexibility of this platform. Highlighting this fact, our replicon-NLC platform has been successfully used for several novel vaccine candidates administered IM [26–30], with initial proof-of-concept studies for IN delivery [8]. In addition to a low toxicity profile when administered IM [30], our NLCs are well positioned for mucosal delivery due to the incorporation of the cationic lipid DOTAP, which can potentially adhere with the many anionic receptors found in the mucosal layer for enhanced delivery and an improved immune response [19,31,32]. However, the adaptation of the replicon-NLC vaccine complex for IN administration requires a more complete understanding of the biophysical and physicochemical properties of the vaccine since properties such as size and charge may affect the mucoadhesive and mucus-penetrative ability of the vaccine [33,34].

As proof of concept, here we report the development and optimization of an H5 hemagglutinin (HA) replicon-NLC vaccine formulation that is equally suitable for IN or IM administration. We then apply this technology to target H5N1 avian influenza virus as a timely proof-of-concept antigen for demonstration of vaccine immunogenicity and efficacy in preclinical animal models. We demonstrate a positive relationship between the nitrogen-to-phosphate (N:P) ratio used to form the replicon-NLC vaccine complex and vaccine immunogenicity, which further enhances our understanding of the physical structure of the vaccine. We also demonstrate that the vaccine is well tolerated in mouse and ferret models and stimulates robust systemic immunity when delivered either IM or IN. Moreover, we establish mucosal immunity after IN but not IM delivery. Importantly, this vaccine formulation was effective in preventing mortality and morbidity in a lethal H5N1 challenge ferret model. Together, this work suggests the utility of this replicon-NLC complex as a potentially powerful and effective replicon vaccine platform that can be adapted for various routes of administration with little to no formulation redevelopment effort required.

## MATERIALS AND METHODS

### NLC formulation production

The NLC formulation was prepared according to an established protocol for research-grade NLC [26,27]. Briefly, squalene (MilliporeSigma), sorbitan monostearate (Span 60; Spectrum Chemicals), 1,2-dioleoyl-3-trimethylammonium-propane (DOTAP; SAFC), and trimyristin (Dynasan 114; MilliporeSigma) were mixed and dissolved by heating to 65°C and sonication in a water bath sonicator to form the oil phase. Separately, polysorbate 80 (Tween 80; MilliporeSigma) was mixed and dissolved in a solution of 10 mM sodium citrate by heating to 65°C with water bath sonication to form the aqueous phase. After both the oil phase and aqueous phase were fully dissolved, the two were rapidly mixed at a rate of 7,000 rpm using a high-speed laboratory emulsifier (Silverson Machines). High-shear homogenization was then carried out on the resultant suspension using a microfluidizer (M-110P, Microfluidics International Corp.). The mixture was processed at 30,000 psi (207 MPa) for 10 discrete microfluidization passes. The final NLC product contained 30 mg/mL DOTAP, 37.5 mg/mL squalene, 2.4 mg/mL trimyristin, 37 mg/mL polysorbate 80, and 37 mg/mL sorbitan monostearate. The hydrodynamic diameter of the NLCs was measured to be 60 ± 25 nm, as determined by dynamic light scattering (DLS; described below), and zeta potential was determined to be 45 ± 5 mV by electrophoretic light scattering (described below). NLCs were terminally filtered using a 0.22 µm polyethersulfone filter and stored at 2 – 8°C in 3 mL glass vials until further use.

### Design of DNA template for replicon production

Template DNA plasmids for the replicon were created by subcloning gene blocks with flanking Gibson assembly overlaps containing a codon-optimized influenza A/Vietnam/1203/2004 H5N1 HA antigen (GenBank ID EU122404.1) into the PflFI/SacII sites in a replicon template plasmid derived from the attenuated TC83 strain of Venezuelan equine encephalitis virus [26,27,35]. The resulting H5-expressing replicon (H5 replicon) template plasmid was sequence-confirmed, amplified in *Escherichia coli,* and linearized by NotI restriction digestion.

### Replicon synthesis and purification

Production and purification of the H5 replicon was performed as described previously [8,27,28]. Briefly, *in vitro* transcription was conducted using the linearized plasmid template and a T7 RNA polymerase-mediated transcription reaction, followed by removal of template DNA by DNase I treatment and addition of cap0 structures using vaccinia capping enzyme. The capped RNA was purified from the reaction components via Akta liquid chromatography using Capto Core 700 multimodal chromatography resin (Cytiva), followed by diafiltration and concentration using tangential flow filtration. The final H5 replicon material was terminally filtered using a 0.22 µm polyethersulfone filter. Purity was assessed by agarose gel electrophoresis, and the replicon was quantified via UV-vis spectrophotometry (NanoDrop, Thermo Fisher Scientific) as well as a commercially available Quant-it RiboGreen RNA assay (Thermo Fisher Scientific). The replicon was stored at -80°C until further use.

### Replicon-NLC complexing

Replicon-NLC vaccine complexes were formed as described previously [8,27,28] with minor modifications. Briefly, complexes were formed at various N:P ratios, defined as the molar charge ratio between the positively charged nitrogen (N) of the lipid DOTAP and the negatively charged phosphate (P) groups of the nucleic acid backbone. Complexes were formed using N:P ratios of 15, 12, 10, 8, 5, 1, and 0.6. Fresh complexes were prepared by mixing 200 µg/mL replicon 1:1 by volume with NLC in a buffer containing 10 mM sodium citrate and 20% w/v sucrose. This resulted in a final isotonic replicon-NLC complex containing 100 µg/mL H5 replicon, 10% w/v sucrose, and 5 mM sodium citrate. The resulting solutions were allowed to incubate on ice for 30 minutes to ensure complete complexing prior to use.

### Size characterization by dynamic light scattering

DLS size measurements were carried out using a Zetasizer Nano ZS instrument (Malvern Panalytical). NLCs or vaccine complexes were diluted 100-fold using nuclease-free water and individually loaded into 40 µL disposable microcuvettes (Malvern Panalytical). The dispersant was set to water at 25°C with a viscosity of 0.8872 cP and a refractive index of 1.330, and the dispersant viscosity was used for the sample viscosity. The automatic attenuation setting was selected, and a backscatter angle of 173° was used. Samples were incubated at 25°C for 30 seconds inside of the instrument prior to collecting measurements. Measurements were recorded in triplicate and averaged, and standard deviation was calculated.

### Zeta potential measurements by electrophoretic light scattering

Zeta potential measurements were carried out using a Zetasizer Nano ZS instrument (Malvern Panalytical). Complexes were diluted 100-fold using nuclease-free water, and 850 µL of each sample was individually loaded into a disposable folded capillary zeta cell (Malvern Panalytical).

Temperature, viscosity, and refractive index settings were the same as used for DLS measurements (see above), and a dielectric constant of 78.5 was used. A Smoluchowski model [36] was used with an F(ka) value of 1.50. The measurement duration was set to automatic, with a minimum of 10 runs and a maximum of 30 runs collected per measurement. A total of five repeat measurements were collected, with a 60-second delay between repeat measurements. The attenuation selection and voltage selection were both set to automatic, and a monomodal analysis model was selected. Samples were incubated at 25°C for 30 seconds inside of the instrument prior to collecting measurements.

### Nanoparticle tracking analysis

Nanoparticle tracking analysis (NTA) was carried out using a NanoSight NS300 instrument equipped with a 532 nm laser (Malvern Panalytical). For the most accurate particle sizing and quantitation, particles should be diluted such that the average number of particles per frame is 20 – 60. To account for how the particle concentration will change as a function of N:P ratio, complexes were diluted with water, a dilution factor of 5000 was used for the N:P range from 15 to 10, and a dilution factor of 2500 was used for the N:P range from 8 to 0.6. Due to the low particle concentration observed at an N:P of 1, a dilution factor of 200 was used for this sample. 1 mL of diluted complex was loaded into a l mL plastic syringe and connected to the instrument feed line via a syringe pump. Samples were flowed through the system for 2 minutes at room temperature using an instrument pump speed setting of 35 arbitrary instrument units prior to capturing five replicate 60-second videos using a camera level setting of 16. Captured videos were then processed using a detection threshold of 6. Histograms, particle size, and particle concentration data were corrected for the dilution factor and averaged, and standard error of the mean was calculated.

### Replicon-NLC integrity and protection assay

Replicon nucleic acid integrity after complexing and protection from RNase-mediated degradation were evaluated using an agarose gel-based method described previously [27]. Briefly, replicon- NLC complexes or control H5 replicon freshly thawed from -80°C were diluted to a replicon nucleic acid concentration of 40 ng/µL and subsequently treated with phenol:chloroform:isoamyl alcohol (25:24:1 v/v; Thermo Fisher Scientific) in a 1:1 ratio by volume. Samples were vortexed and subsequently centrifuged at 17,000 x *g* for 15 minutes to separate the organic and aqueous phases. The aqueous phase was mixed 1:1 by volume with glyoxal load dye (Invitrogen), heated at 50°C for 20 minutes to denature the replicon, then cooled at room temperature for 5 minutes prior to loading on an agarose gel.

Separately, complexes or control H5 replicon were diluted to 40 ng/µL and treated with RNase A (Thermo Fisher Scientific) in a 1:40 RNase A:nucleic acid ratio for 30 minutes at room temperature. RNase A reactions were then immediately quenched by the addition of recombinant Proteinase K (Thermo Scientific) at a ratio of 1:100 RNaseA:Proteinase K and heated for 10 minutes at 55°C. The quenched reaction was cooled to room temperature, and replicon nucleic acid was then extracted using phenol:chloroform:isoamyl alcohol, vortexing, and ultracentrifugation as described above. The aqueous phase was mixed 1:1 by volume with glyoxal load dye (Invitrogen) and heated at 50°C for 20 minutes, then cooled back down to room temperature before loading on an agarose gel.

For each sample and control, 300 ng of replicon nucleic acid was loaded onto a 1% w/v denaturing agarose gel. Gels were run at 120 V for 45 minutes at room temperature. Gels were imaged using an ethidium bromide protocol on a ChemiDoc MP imaging system (Bio-Rad Laboratories). Gel densitometry analysis was performed using Image Lab Software version 6.1.0 (Bio-Rad Laboratories). Relative nucleic acid integrity of extracted and RNase-challenged replicon samples was determined by comparing band intensities with that of the replicon loading control (uncomplexed replicon that was subjected to the extraction process and not treated with RNase A).

### Cryogenic transmission electron microscopy (Cryo-TEM) of NLCs

Cryo-TEM was performed at NanoImaging Services (San Diego, CA). Replicon-NLC complexes were formed with 100 ng/µL replicon at N:P ratios of 15 and 0.6 as described above. The samples were then diluted with formulation buffer to an estimated particle concentration of ∼10^12^ to 10^13^ particles/mL. A 3 μL drop of NLC formulation was applied to a 300-mesh 2/1 C-flat copper grid (Protochips) that had been plasma-cleaned for 10 seconds using a Solarus plasma cleaner (Gatan Inc.). The grid was blotted for 6 seconds at 4°C and 100% humidity to remove excess liquid and form a thin aqueous film and was subsequently frozen by submersion into liquid ethane cooled by liquid nitrogen using a Vitrobot automated plunger (Thermo Fisher Scientific). Vitreous ice grids were clipped into cartridges, transferred into a cassette, and then into the Glacios autoloader, all while maintaining the grids at cryogenic temperature (below -170°C). Data collection was performed using a Glacios Cryo-Transmission Electron Microscope (Thermo Fisher Scientific) operated at 200 kV and equipped with a Falcon 3 direct electron detector. Automated data collection was carried out using Leginon software [37,38], where high magnification movies were acquired by selecting targets at a lower magnification. Images of each grid were acquired at multiple scales to assess the overall distribution of the specimen. After identifying potentially suitable target areas for imaging at lower magnifications, high magnification images were acquired at nominal magnifications of 150,000x (0.095 nm/pixel), 73,000x (0.201 nm/pixel), and 28,000x (0.519 nm/pixel). Images were acquired at a nominal underfocus of -5.5 μm to -2.0 μm and electron doses of ∼10-25 e-/Å^2^.

### *In vitro* cell uptake assay

To assess the correlation between N:P ratio and cellular uptake, Vero cells (American Type Culture Collection #CCL-81) were transfected with a green fluorescent protein (GFP)-expressing replicon complexed with NLC at different N:P ratios. Vero cells were cultured in 6-well plates to 80% confluency (approximately 1 million cells) the day of transfection. The replicon and NLC were complexed at varying N:P ratios from 0.6 to 15 at a concentration of 1 µg/mL in vaccine formulation buffer (described above). Vero cell monolayers were washed two times with warm phosphate-buffered saline (PBS), overlaid with 1 mL of each formulation in triplicate, and left to incubate at 37°C and 5% CO2 for 4 hours. After incubation, cells were overlaid with 2 mL Dulbecco’s Modified Eagle Medium + 10% fetal bovine serum (FBS), then imaged using an ImageXpress Pico imager (Molecular Devices) to detect GFP expression 24 hours post-transfection, using an exposure time of 200 milliseconds for the 488-nm channel.

### Mouse studies

Mouse studies were performed in compliance with all applicable sections of the Final Rules of the Animal Welfare Act regulations (9 CFR Parts 1, 2, and 3) and the *Guide for the Care and Use of Laboratory Animals* [39]; and under the oversight of the Bloodworks Northwest Research Institute (Seattle, WA) Institutional Animal Care and Use Committee (IACUC), protocol #5389-01.

C57BL/6J mice from The Jackson Laboratory, between 6 and 8 weeks of age at study onset, were used for all mouse studies. All-female mice were used in the initial formulation screen (**Figure 5**) to maximize statistical power, while equal numbers of male and female mice were used in the detailed immunogenicity study (**Figure 6**). Each mouse was administered vaccine either IN or IM at 5 µg replicon per mouse as indicated at day 0 (prime vaccination) and day 21 (boost vaccination) of the study. Mice were vaccinated with a replicon expressing the H5 antigen or secreted embryonic alkaline phosphatase (SEAP; negative control). Each mouse was individually ear-tagged and tracked across the entire study. Blood samples were collected by the retro-orbital route every 7 days on isoflurane-sedated mice, and terminal bleeds were collected by cardiac puncture after euthanasia at study end. Post-prime and post-boost serum was analyzed for H5-binding IgG titer by ELISA. At study end, spleens, lungs, bronchoalveolar lavage (BAL), and/or bone marrow were collected and stored appropriately on ice for additional processing and use in a variety of downstream immunoassays to quantify the immune response to vaccine regimen.

Mice were euthanized early under the following conditions: 20% weight loss relative to starting bodyweight, lack of mobility, lethargy, and/or hunched back that did not resolve. Mice were euthanized using CO2 euthanasia as the primary method, and cervical dislocation as the secondary method, in accordance with the recommendation of the Panel on Euthanasia of the American Veterinary Medical Association. Death was confirmed by absence of corneal reflex.

### Peptide stimulation and intracellular cytokine staining-flow cytometry

Spleens and lungs were processed to harvest single-cell suspensions as previously described [8,28,35]. Stimulation of spleen and lung single-cell suspensions was performed as previously described [28,35]. Briefly, RPMI 1640 + 10% FBS + beta-mercaptoethanol were prepared containing anti-CD28 antibody and brefeldin A. Cells were stimulated with one of three stimulation treatments: (1) 0.26% dimethyl sulfoxide (DMSO) as a negative stimulation control, (2) 1 μg/mL H5 peptide pool (JPT Peptide Technologies #PM-INFA-HAIndo; pool derived from A/Indonesia/CDC835/2006(H5N1), which has a 96.64% sequence homology with the H5 HA from A/Vietnam/1203/2004 used in these studies) in an equivalent amount of DMSO, or (3) 10 μg/well of phorbol myristate acetate-ionomycin solution as a positive stimulation control. After 6 hours of incubation at 37°C and 5% CO2, samples were washed with 1X PBS, centrifuged again, and stained for flow cytometry [28]. Cells were first stained for viability with Zombie Green (BioLegend), followed by anti-CD16/32 (Invitrogen #14-0161-86) to block Fc receptors. All samples were surface stained for CD4 (APC-Cy7, BD #565650), CD8α (BV510, BioLegend #100752), and CD44 (PE-CF594, BD #562464) with lungs additionally stained for CD69 (PE, BD #553237) and CD103 (BV711, BioLegend #121435). Cells were fixed and permeabilized using a Fixation/Permeabilization Kit (BD Biosciences #554714) followed by intracellular staining for IFN-γ (PE-Cy7, BioLegend #505826), TNF-α (BV421, BioLegend #506328), IL-2 (PE-Cy5, BioLegend #503824), IL-5 (APC, BioLegend #504306), and IL-17A (AF700, BioLegend #506914). Samples were run on a CytoFLEX flow cytometer (Beckman Coulter) and analyzed with FlowJo software (BD Biosciences).

### Bone marrow ELISpot assay

Antibody-secreting cells in bone marrow were assessed by ELISpot assay as previously described [40]. Briefly, plates were coated with recombinant H5 A/Vietnam/1194/2004 antigen (Sino Biological). Single-cell suspensions of bone marrow samples collected within the last 12 hours were seeded at 1 x 10^6^ cells per well and serially diluted and then probed with a goat anti-mouse IgG horseradish peroxidase conjugate antibody (SouthernBiotech #1031-05). 3-amino-9- ethylcarbazole (ACE) substrate kit (Vector Laboratories) was used as the colorimetric substrate for the ELISpot development. Colored spots were enumerated using an automated ELISpot reader (CTL Analyzer, Cellular Limited Technology). Data were analyzed using ImmunoSpot software (Cellular Limited Technology).

### Ferret efficacy study

Ferret studies were performed in accordance with national and institutional guidelines for animal care of laboratory animals and approved by the Colorado State University IACUC, protocol #4228, as previously described [35]. Fitch ferrets (*Mustela putoris fero*) were purchased from Triple F Farms at approximately 3 months of age and confirmed to be serologically negative for influenza virus by microneutralization testing with H5N1-PR8 recombinant influenza virus prior to initiating immunization. All ferrets were castrated or spayed prior to the study. Vaccination with replicon- NLC complexes consisted of a prime dose of 10 µg replicon given either IM or IN to a cohort of 6 ferrets per group (3 male, 3 female). A subsequent boost dose of 10 µg replicon was delivered IN 21 days post-prime. An alum-adjuvanted inactivated whole virion H5N1 vaccine (“Baxter;” BEI Resources, NIAID, NIH: A/H5N1 Influenza Vaccine, Inactivated Whole Virion (A/Vietnam/1203/2004), Vero-Cell Derived, Adjuvanted, 15 Micrograms HA, NR-12143) was used as a positive control, while SEAP replicon-NLC was used as a negative control.

Ferrets were observed daily for 3 days after vaccination for any signs of adverse reactions, and observed daily for symptoms and clinical scores after challenge until necropsy. Clinical scores were defined as 0 = normal; 1 = possibly less active than normal OR not grooming as normal (questionably sick or mild illness); 2 = less active than normal when viewed in cage but appears nearly normal and responsive when handled (sick); 3 = reluctance to rise when viewed in cage and not normally responsive when handled by humans OR weight loss up to 15% of pre-challenge value; 4 = failure to move when stimulated by humans OR collapse when moving OR recumbency OR weight loss >20% of pre-challenge value OR manifestation of any neurologic signs (ataxia, head tilt, etc.) OR body temperature < 100°F; and 5 = found dead. Serum samples were taken pre-challenge, and animal clinical scores and nasal wash viral loads were assessed after challenge.

Challenge was conducted as previously described [41,42]. Ferrets were lightly anesthetized with ketamine-xylazine, and 0.5 mL (0.25 mL/nostril) containing 10^6^ PFU of influenza A/Vietnam/1203/2004 H5N1 virus was instilled IN. Following infection, animals were observed and temperatures were recorded at least once daily. The animals were weighed at the time of challenge and every day thereafter. Nasal washes were collected under ketamine-xylazine sedation on Days 1, 3, and 5 post-challenge, and viral titers were quantified by plaque assay on Madin-Darby canine kidney cells. Any animal that displayed severe clinical signs or that lost > 25% of pre-challenge weight was euthanized at that time, and survivors were euthanized on Day 14 post-challenge.

### Quantification of IgG and IgA antigen-binding titers by ELISA

Antigen-binding IgG in serum and IgA in BAL samples were quantified by ELISA as previously described [28,35]. Plates were coated with influenza H5(H5N1) (A/Vietnam/1203/2004) (Immune Technology #IT-003-0051p) and blocked for at least 1 hour. Mouse serum samples were diluted to create a 14-point dilution curve starting at 1:40 for serum or 1:4 for BAL. A 12-point dilution curve starting at 1:160 was used for ferret serum. Negative controls included naïve C57BL/6J mouse serum or pre-immunized fitch ferret serum and were diluted identically to the samples. Positive controls included an anti-HA antibody (HA from A/Vietnam/1203/04 influenza virus; Rockland Immunochemicals #200-301-976) for mice or inactivated post-H5N1 infection pooled serum for ferrets. Serially diluted samples were transferred onto coated plates and incubated for 1 hour. H5- binding IgG in serum was detected using the Anti-Mouse IgG (Fc specific)-Alkaline Phosphatase antibody (Sigma-Aldrich #A2429) or an anti-ferret IgG (H&L specific)-horseradish peroxidase antibody (Abcam #ab112770), both diluted 1:4000 in blocking buffer. H5-binding IgA in BAL samples was detected using the Anti-Mouse IgA (α-chain specific)-Alkaline Phosphatase antibody (Sigma-Aldrich #A4937), diluted 1:1000 in blocking buffer. Plates were then developed with appropriate substrate and read spectrophotometrically at 405 nm. Sample concentrations were interpolated off the linear region of each sample dilution curve using the standard curve for absolute quantification of IgG titers in mouse serum or by calculating endpoint titers from a four-parameter logistic curve fit for IgG titers in ferret serum and IgA in mouse BAL samples. Lower limits of detection (LLODs) were calculated by taking the slope of the linear portion of the four-parameter logistic curve and dividing that by three times the standard deviation of the negative control samples.

### Statistical analysis

Serum IgG, bone marrow IgG, nasal wash viral titers, and antigen-specific T cell responses were first log- or log(y+1)-transformed and then assessed by one-way ANOVA or mixed-effects analysis with Dunnett’s or Tukey’s multiple comparisons test. BAL IgA was assessed by Kruskal-Wallis with Dunn’s multiple comparisons test. All statistical calculations were performed using GraphPad Prism 10.0 software.

## RESULTS

### N:P ratio dictates vaccine structure and replicon protection from nuclease-mediated degradation

The N:P ratio is a key parameter that influences the physicochemical characteristics of the replicon-NLC vaccine complex [43,44], which may, in turn, affect how well the vaccine binds to and traverses the nasal mucosa upon IN administration. Therefore, the N:P ratio of the vaccine complex must be optimized to achieve appropriate physicochemical properties that are conducive to vaccine uptake and antigen expression when administered IN. More specifically, sufficient vaccine mucoadhesion is necessary to prevent rapid drug clearance prior to intracellular uptake. Mucoadhesiveness is frequently dictated by the surface properties of the particle, i.e., the vaccine zeta potential. Cationic nanoparticles typically have high mucoadhesiveness due to electrostatic interactions with negatively charged mucin glycoproteins [33], making the intrinsically cationic NLC platform well suited for mucosal administration. However, due to the structure of the replicon-NLC complexes wherein the replicon nucleic acid is not encapsulated within the NLC but rather electrostatically bound to the NLC surface, we hypothesize that the vaccine N:P ratio (the ratio of positively charged NLC amines (N) to negatively charged phosphates (P) in the replicon nucleic acid backbone) will influence the structural organization of the formed complexes, and thereby their measured zeta potential and potentially the degree of replicon protection from nuclease-mediated degradation. We therefore first evaluated the effect of N:P ratio on the apparent zeta potential of the H5 replicon vaccine complex.

NLCs were complexed with the H5 replicon at N:P ratios ranging from 0.6 to 15, and the apparent zeta potential of the formed complexes was subsequently measured. It was observed that vaccine complexes had a positive and constant zeta potential of +14 mV when formed at N:P ratios ranging from an N:P of 15 down to an N:P of 5 (**Figure 1A, Supplementary Table 1**). Further decreasing the N:P ratio to an N:P of 1 and below resulted in a sharp decrease in the apparent zeta potential from +14 mV to -37 mV, suggesting that a threshold had been reached whereby the cationic charge of the NLC was overcome by a saturating anionic charge of the replicon that is electrostatically bound to the NLC surface. This was in agreement with agarose gel assessment of whole replicon- NLC vaccine complexes, which show the presence of excess replicon (in the form of an ethidium bromide-stained smear in the gel lane) at N:P ratios less than or equal to 1, indicating that the replicon has fully saturated the NLCs at these N:P ratios (**Supplementary** Figure 1); at higher N:P ratios, no excess replicon was observed, indicating complete association with the NLC surface. Further decreasing the N:P ratio to an N:P of 0.6 (empirically determined to be the saturating N:P ratio based on the sudden appearance of a distinct replicon band on an agarose gel) further decreased the surface charge to -54 mV (**Figure 1A, Supplementary Table 1**), reflecting the increased presence of replicon nucleic acid in the complex and decreased exposure of the cationic NLC surfaces.

**Figure 1.**
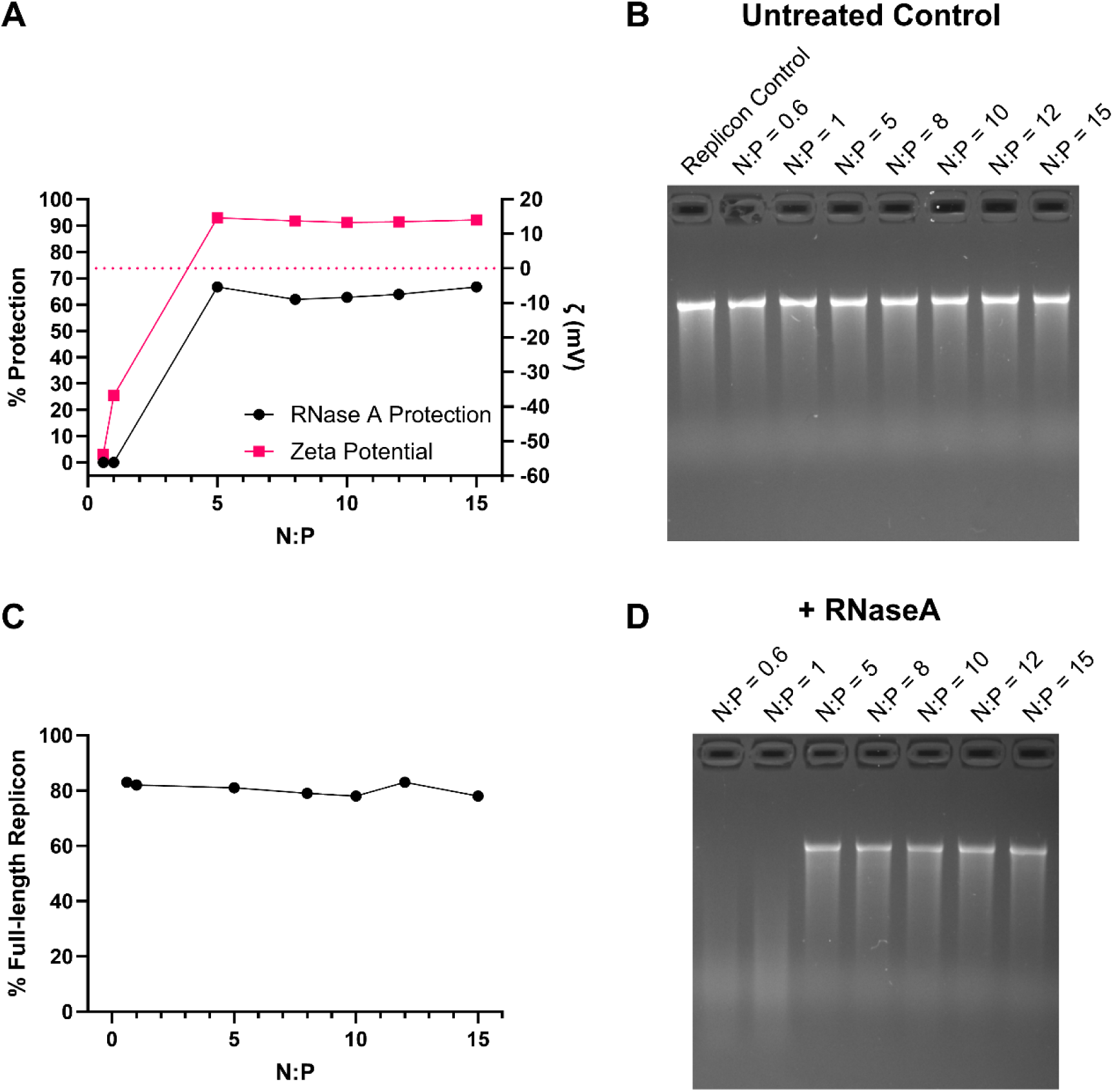
Assessment of full-length H5 replicon nucleic acid protection at varying N:P ratios. (A) Co-plot of % protection of the H5 replicon from enzymatic degradation at varying N:P ratios (black) and apparent zeta potential (pink). The % protection is defined as the agarose gel band intensity of the RNase A-treated H5 replicon relative to the untreated H5 replicon. (B) Agarose gel of replicon nucleic acid after complexation with NLCs at varying N:P ratios and subsequent organic extraction. (C) The percentage of full-length replicon construct from the agarose gel in panel B as a function of the N:P ratio, relative to a freshly prepared and non-complexed replicon control band. (D) Agarose gel of replicon nucleic acid after complexation with NLCs at varying N:P ratios, followed by enzymatic challenge with RNase A and subsequent organic extraction.

We next assessed the effect of the N:P ratio on protection of the H5 replicon construct from nuclease-mediated degradation by briefly treating complexes formed at various N:P ratios with RNase A, followed by organic extraction of the nucleic acid and subsequent agarose gel assessment. At all N:P ratios tested in the absence of RNase A, the replicon construct was well maintained after complexing with NLC, relative to a freshly prepared pure replicon control (**Figure 1B-C**). Upon challenge with RNase A, at N:P ratios ≥ 5, a high and consistent degree of H5 replicon protection from nuclease-mediated degradation was observed (**Figure 1D**). Interestingly, the complete degradation of full-length replicon nucleic acid was observed at N:P ratios ≤ 1. This appeared to coincide with a change in the apparent zeta potential from positive to negative values (**Figure 1A**).

To better understand the structural implications of the relationship between N:P ratio, complex zeta potential, and replicon protection, cryo-TEM was performed on vaccine complexes formed at an N:P of 15 and an N:P of 0.6, which represent the two extremes of zeta potential and replicon protection. TEM imaging revealed contrasting structures formed at these two N:P ratios: at an N:P of 15, NLC structures were ubiquitous throughout the image, and replicon structures were not readily apparent, suggesting the tight compaction of the replicon nucleic acid within multiple NLC molecules (**Figure 2A**). We also observed the presence of rod-like structures, which may be aggregates formed from a small excess of lipids that may be present at high N:P ratios. In contrast, at an N:P of 0.6, we observed a large amount of replicon nucleic acid density surrounding sparse populations of NLCs and the absence of these rod-like structures (**Figure 2B**). This variability in the observed replicon density at different N:P ratios may explain the measured zeta potential and differences in replicon protection (see Discussion). Taken together, these data suggest that an N:P ratio ≥ 5 will be necessary to maintain a positive zeta potential suitable for replicon-NLC vaccine complex mucoadhesion as well as provide adequate replicon protection to achieve efficient IN delivery of the vaccine.

**Figure 2.**
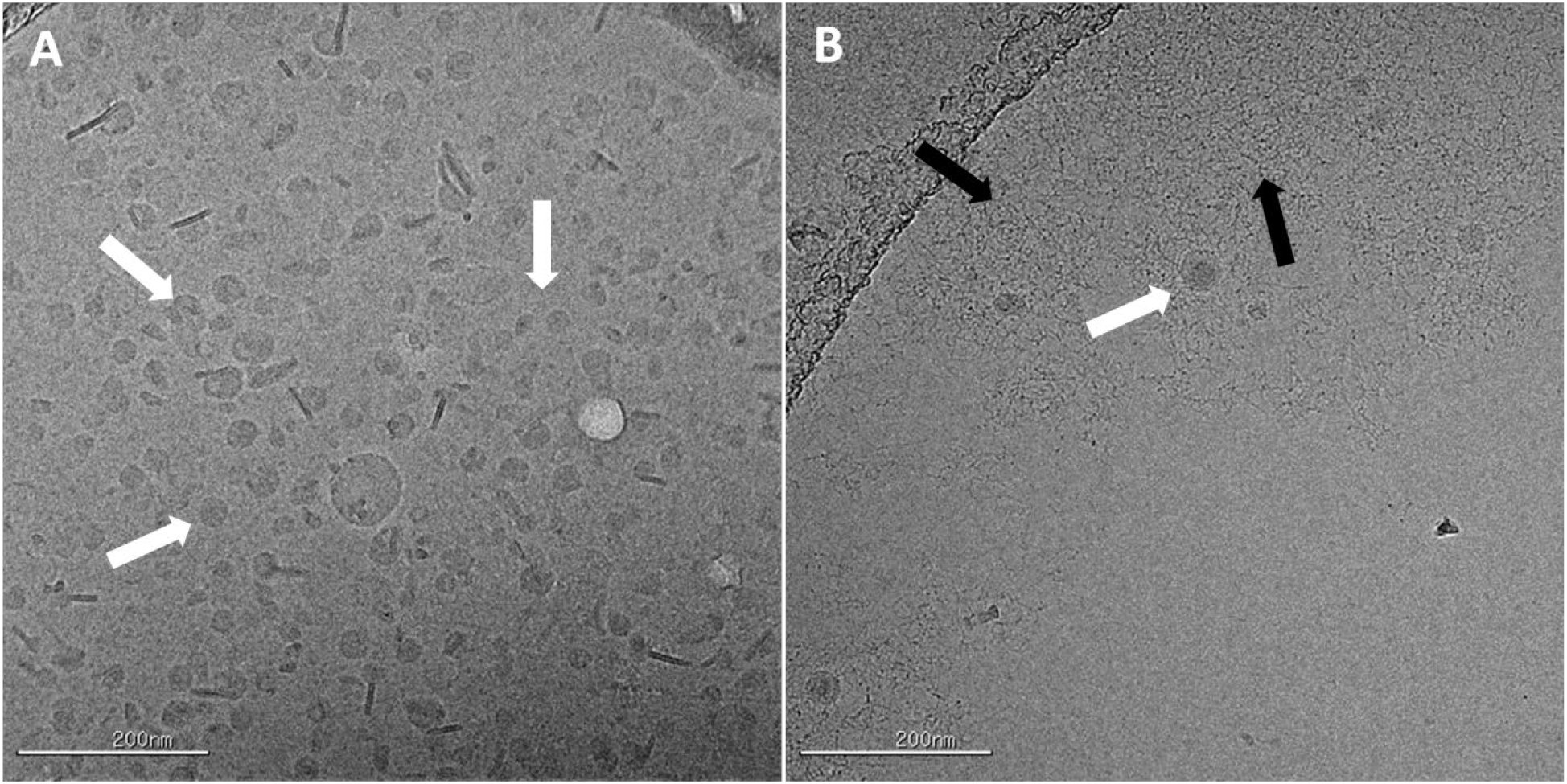
Cryo-TEM images of replicon-NLC complexes. (A) Replicon-NLC complexes formed at N:P of 15. (B) Replicon-NLC complexes formed at N:P of 0.6. White arrows = individual NLC molecules; black arrows = regions of high replicon density. Scale bar = 200 nm.

### Higher N:P ratios promote H5 replicon vaccine uptake by increasing the total nanoparticle concentration of the active complex

Without adequate mucopenetration to traverse the complex nasal mucosa, an IN vaccine will not elicit an efficient immune response prior to mucosal clearance. The mucopenetrative ability of a nanoparticle appears to be related to its size, although this is likely dependent on the specific nanoparticle formulation [32–34]. We therefore decided to test the effects of the N:P ratio on the size of the H5 replicon-NLC complex. NLCs were complexed with the H5 replicon at N:P ratios ranging from 0.6 to 15, and the resulting size of the replicon-NLC complexes was characterized by DLS. DLS measurements revealed a polydisperse intensity size distribution that gradually shifted towards larger particle sizes as the N:P ratio was decreased (**Figure 3A**). At an N:P of 1, a sudden large increase in the Z-average diameter was observed, indicating the likely formation of very large aggregate species > 1 µm in diameter (**Figure 3B, Supplementary Table 2**).

**Figure 3.**
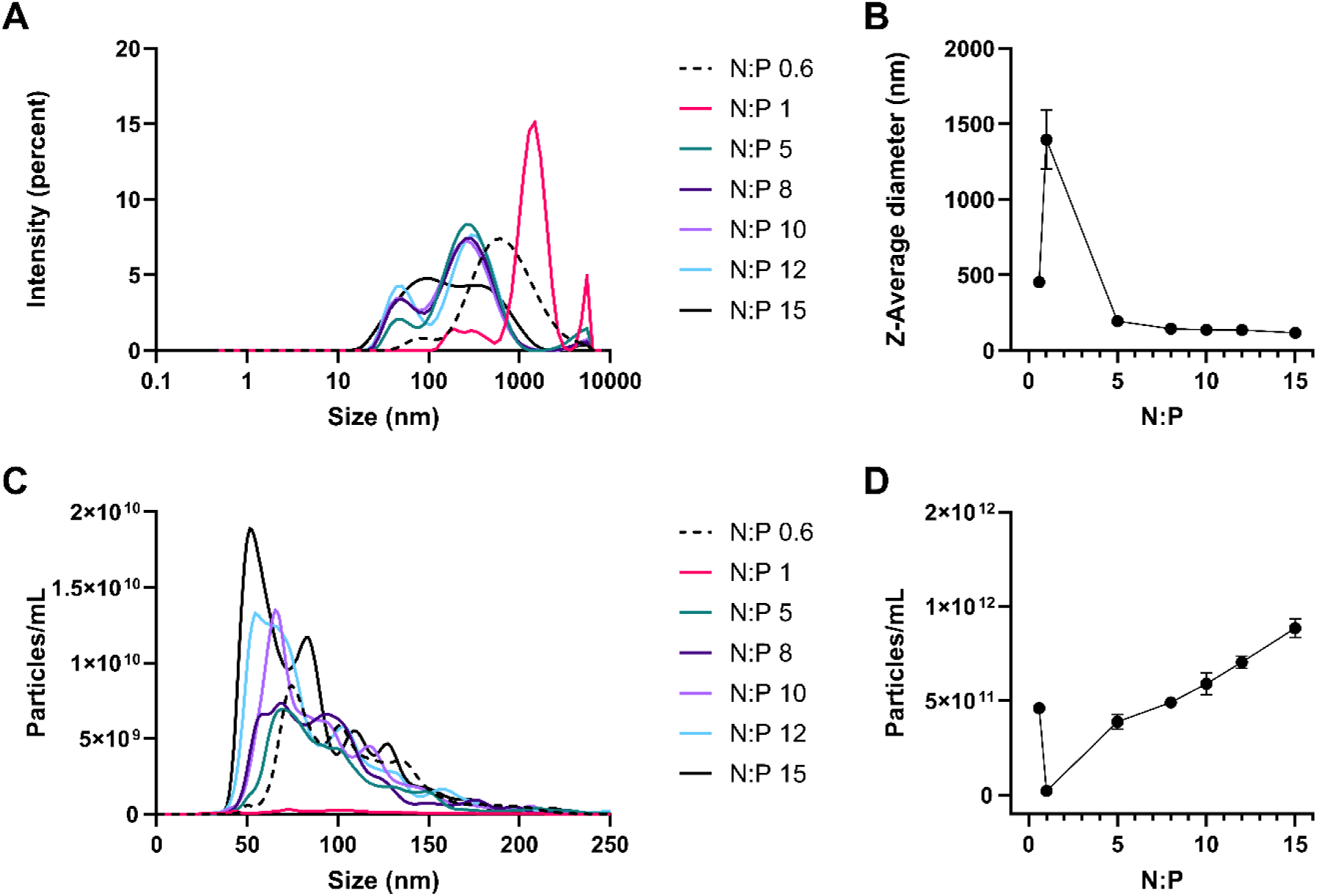
Nanoparticle size and concentration characterization of H5 replicon-NLC complexes at different N:P ratios. (A) DLS intensity size distribution of vaccine complexes formed at varying N:P ratios. Distributions are the average of *n* = 3 repeat measurements. (B) Z-average diameter of H5 replicon-NLC complexes as a function of N:P ratio. Data are presented as the mean ± standard deviation (*n* = 3). (C) NTA size distribution of vaccine complexes formed at varying N:P ratios. Distributions are the average of *n* = 5 repeat measurements. (D) NTA particle concentration measurements as a function of N:P ratio. Data are presented as the mean ± standard error (*n* = 5).

Interestingly, further decreasing the N:P ratio to 0.6 resulted in a shift in the size distribution and Z-average to a size smaller than that observed at an N:P of 1. However, this Z-average particle size was still notably larger than the particle sizes observed for N:P ratios ≥ 5 (**Figure 3A-B**), suggesting that a size maximum is being approached as the N:P ratio approaches a value of 1. Overall, these data suggest that the vaccine complex nanoparticle size is loosely correlated with the N:P ratio in a N:P range of 5 – 15, and that this correlation is lost at N:P ratios ≤ 1, indicating increased complexity in complex formation.

Due to the intrinsically polydisperse nature of the replicon-NLC vaccine complex, we recognized that DLS alone is not sufficient to fully understand the effects of N:P ratio on complex size. Thus, NTA, an orthogonal sizing technique, was next used to investigate the vaccine complex size distribution as a function of N:P ratio. Using capture settings optimized for detecting replicon- NLC complexes, NTA analysis revealed that the nanoparticle size distribution of the vaccine complexes spanned a consistent size range (approximately 40 – 250 nm in diameter) regardless of the N:P ratio used (**Figure 3C**). It was observed that as the N:P ratio was decreased, the total nanoparticle concentration of the vaccine complex was also decreased, until an apparent minimum was reached at an N:P of 1 (**Figure 3D**). Further decreasing the N:P ratio from an N:P of 1 to an N:P of 0.6 resulted in a recovery of nanoparticle concentration. This was consistent with DLS results where the Z-average size reached an apparent maximum at an N:P of 1 and decreased when the N:P ratio was further decreased to an N:P of 0.6 and would suggest that the apparent size differences observed by DLS are likely a function of the total nanoparticle concentration of the replicon-NLC vaccine complex: as the total nanoparticle concentration is increased, the intensity of the scattered light detected during DLS measurements is also increased, resulting in an apparent shift to increasingly larger sizes in the intensity size distribution. Though, it should be noted that NTA cannot reliably detect large aggregate species greater than 1 µm in diameter that were apparent in the DLS size distributions at low N:P ratios [45].

With clear effects on the biophysical properties of zeta potential, protection from nuclease-mediated degradation, and complex particle concentration, we next evaluated how N:P ratio affects the transfection efficiency of the vaccine. Vero cells were transfected with GFP-expressing replicon-NLC complexes formed at various N:P ratios at a 1 µg dose replicon per approximately 1 million cells, and the mean fluorescence intensity was measured. As expected, the degree of GFP expression increased at increasing N:P ratios, indicating increased vaccine uptake by the cells (**Figure 4**). Interestingly, GFP expression was also observed at N:P ratios ≤ 1, albeit to a much lower degree than vaccine complexes formed at N:P ratios ≥ 5, and despite having an apparently negative average zeta potential observed at these ratios (**Figure 4B-C**). This suggested the presence of at least a small amount of nanoparticle species able to transfect cells even at low N:P ratios. Regardless, the N:P ratio appeared to dictate the total nanoparticle concentration of an active NLC vaccine complex that was able to transfect cells for expression of the replicon cargo, with higher N:P ratios resulting in increased replicon uptake and expression. The observation that the necessary N:P ratio is formulation dependent and dictates the degree of intracellular uptake, with more efficient uptake occurring with cationic particles, was previously noted by others for various nanoparticle systems [47,48]. For this system, an N:P ratio ≥ 5 appeared to be necessary to achieve a high enough particle concentration of the active vaccine complex for efficient intracellular uptake and expression of the replicon.

**Figure 4.**
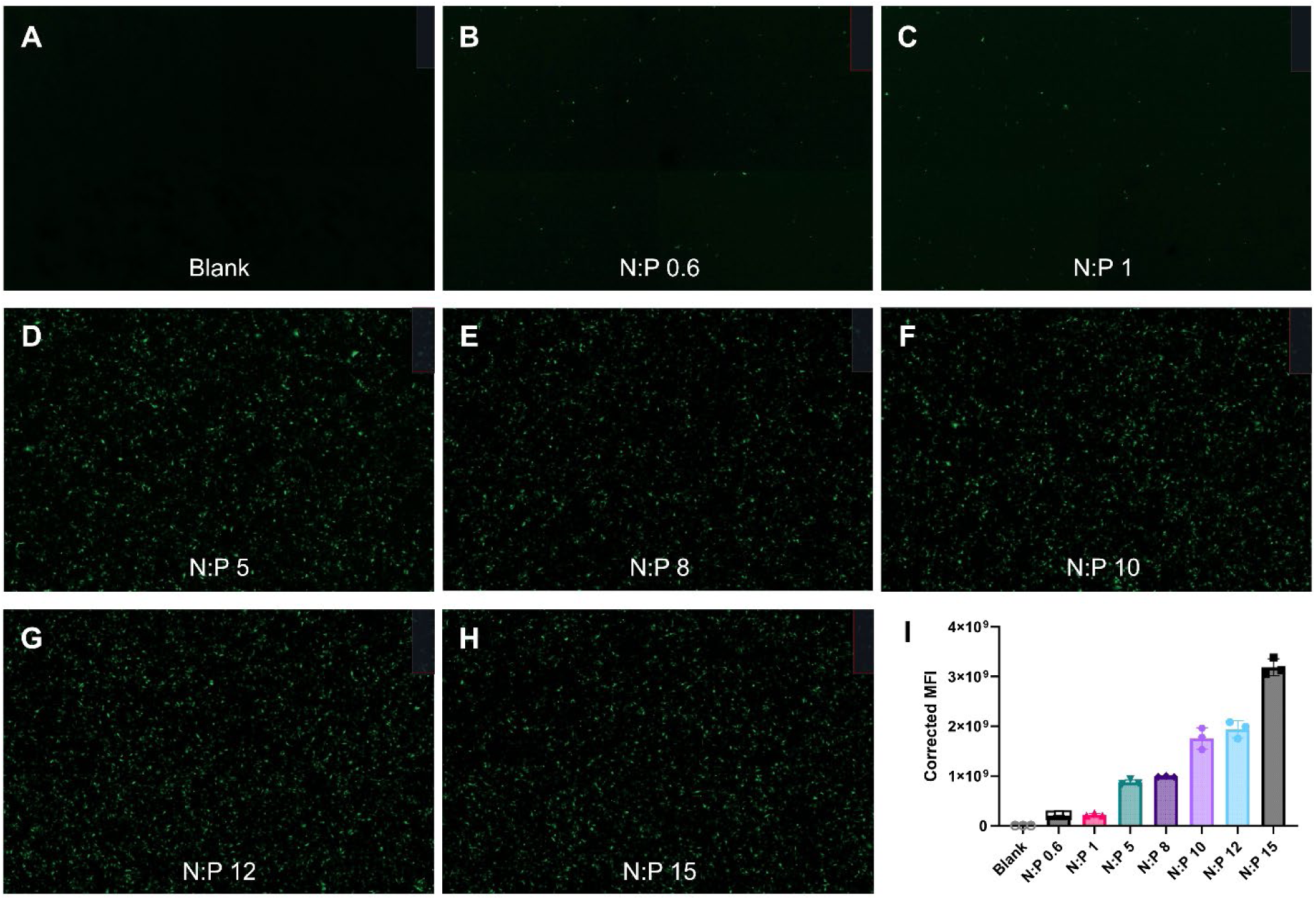
Cellular uptake of replicon-NLC complexes formulated at different N:P ratios. (A-H). Vero cells transfected with GFP-expressing replicon-NLC complexes formed at various N:P ratios. (I) Mean fluorescence intensity (MFI) from triplicate transfected well images. Data points are presented as mean ± standard deviation (*n* = 3 repeat measurements).

### A minimum N:P ratio threshold determines intranasal vaccine immunogenicity

To explore how the N:P ratio affects vaccine-induced immunogenicity *in vivo* following IN administration, C57BL/6 mice were vaccinated IN with a 5 µg dose of H5 replicon complexed with NLC at increasing N:P ratios. As a negative control for H5 immunogenicity, a group of mice was vaccinated IN with a 5 µg dose of a replicon expressing SEAP, complexed with NLCs at an N:P of 15, which is not expected to generate an H5-reactive immune response. As a separate positive control, mice were vaccinated IM with a 5 µg dose of H5 replicon complexed with NLC at an N:P of 15 [8]. Mice received a prime dose on day 0 through either IN or IM administration, and a second dose 21 days after the prime dose via the same route of administration. Serum samples were collected 42 days post-prime dosing to assess humoral responses to vaccination, and bone marrow and spleen were harvested 42 days post-prime dosing to assess humoral and cellular responses, respectively (**Figure 5A**).

**Figure 5.**
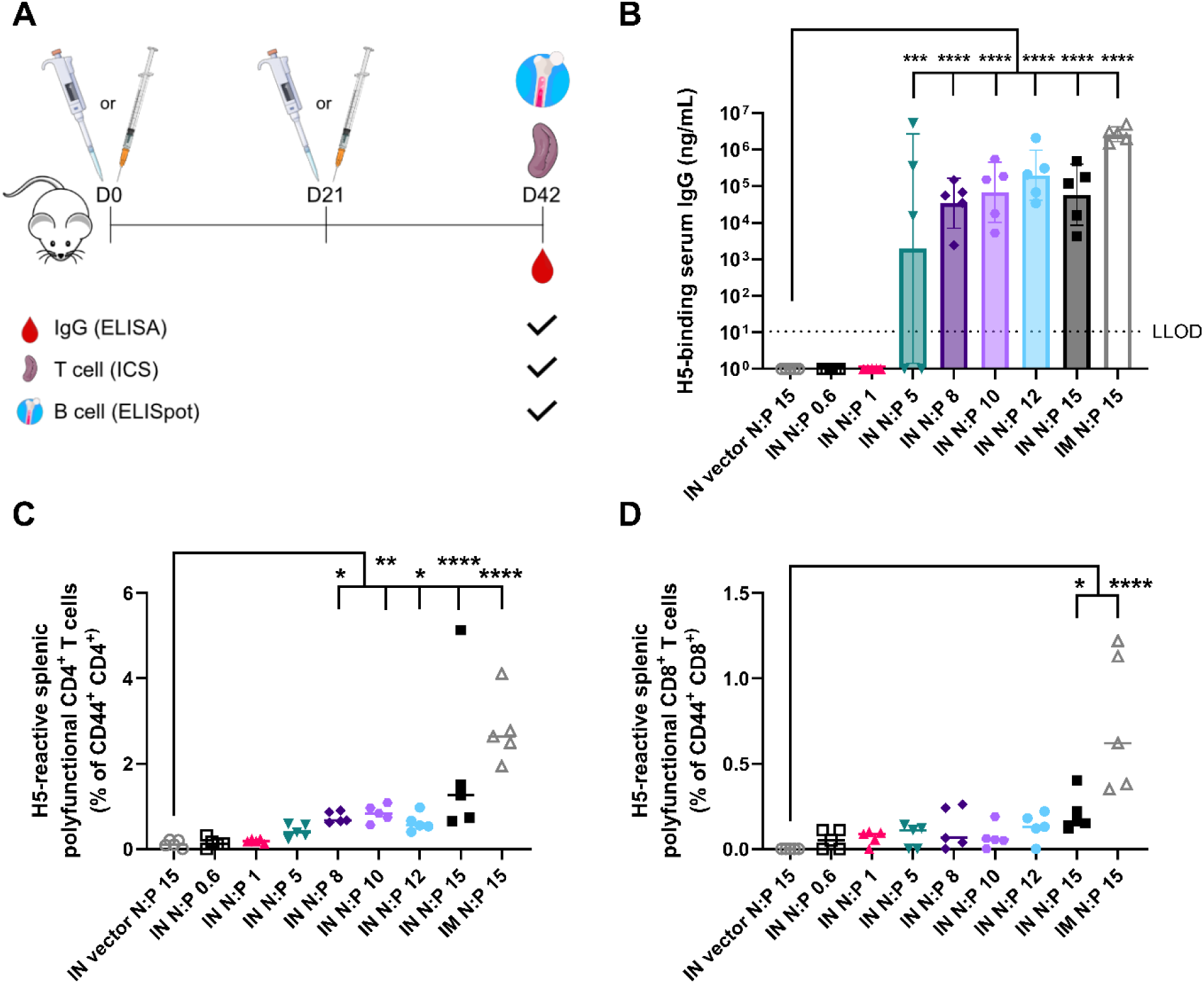
Immunogenicity of 5 µg H5 replicon-NLC vaccine complexes as a function of N:P ratio. A complex formed at N:P of 15 using a SEAP-expressing replicon was used as a negative vector control. *n* = 5 female mice per group. (A) Study design. Bone marrow ELISpot shown in **Supplementary** Figure 2. (B) Influenza H5-binding serum IgG titers in mice dosed with vaccine complexes formed at varying N:P ratios post-boost. Data were log-transformed and displayed as the geometric mean ± geometric standard deviation. Data were analyzed using ordinary one-way ANOVA with Dunnett’s multiple comparisons test. (C-D) ICS flow cytometric analysis of H5-reactive polyfunctional (IFNγ^+^ IL-2^+^ TNFα^+^) splenic (C) CD4^+^ T cells and (D) CD8^+^ T cells in mice dosed with vaccine complexes formed at various N:P ratios. Horizontal lines show median. Data were log(y+1)-transformed and analyzed using one-way ANOVA with Tukey’s multiple comparisons test. * *p* < 0.05, ** *p* < 0.01, *** *p* < 0.001, **** *p* < 0.0001. IM = intramuscular, IN = intranasal, and LLOD = lower limit of detection.

The effect of the N:P ratio on vaccine induction of serum H5-binding IgG titers was evaluated post-boost dose by ELISA. Mirroring what was observed *in vitro*, animals vaccinated IN with the H5 replicon at N:P ratios ranging from 5 to 15 generated IgG titers that were statistically significant over a SEAP replicon vector control (**Figure 5B**). In contrast, H5-binding IgG titers were undetectable in animals vaccinated with H5 replicon-NLC complexes at N:P ratios of 0.6 or 1. Interestingly, in two independent experiments, we observed an intermediate response in animals vaccinated at an N:P of 5 with undetectable titers in some animals and strong titers in others. This trend in vaccine immunogenicity was confirmed by ELISpot analysis of H5-specific IgG-secreting bone marrow cells (**Supplementary** Figure 2). Together, these data suggest that an N:P of 5 is on the threshold between functional and non-functional vaccine formulations when delivered IN.

To evaluate how the N:P ratio affects the cellular immune response to vaccination, we stimulated post-boost splenocytes with overlapping H5 peptides and used intracellular cytokine staining (ICS) and flow cytometry to quantify H5-reactive polyfunctional (IFNγ^+^ IL-2^+^ TNFα^+^) T cells (the gating strategy is described in **Supplementary** Figure 3). Splenic polyfunctional CD4^+^ T cells were detectable for all IN groups dosed with the H5 replicon at an N:P ratio ≥ 5, and similar to ELISA results for H5-binding IgG titers, vaccine complexes formed at an N:P ratio ≥ 8 induced statistically significant populations compared with those induced by a SEAP replicon vector control (**Figure 5C**). Polyfunctional CD8^+^ T cell counts for IN dosed mice were somewhat elevated over those seen in mice dosed with the SEAP control complex (**Figure 5D**). However, this was not statistically significant except for mice dosed with an N:P of 15, and this study was likely underpowered to detect meaningful differences in the polyfunctional CD8^+^ T cell population. All IN dosed mice showed weaker induction of CD4^+^ and CD8^+^ T cells compared to the IM dosed control group at N:P of 15, suggesting that IM dosing of the vaccine may induce a more potent systemic T cell response than is possible with IN vaccination alone. While the available data indicate minimal toxicity of the lipid DOTAP in the NLCs [30], DOTAP’s safety profile after IN administration is not yet widely established; therefore, all subsequent *in vivo* dosing studies were carried out using a vaccine N:P of 10 to achieve a minimal effective NLC dose and to allow for the detection of any observed differences between experimental groups.

### Intranasal replicon-NLC vaccination uniquely stimulates mucosal immune responses, with balanced systemic-mucosal immunity stimulated by heterologous IM/IN immunization

As discussed above, a major advantage of IN vaccines is their potential to stimulate mucosal immune responses that may aid in protection against respiratory virus infections. Additionally, previous vaccine dosing studies indicate that heterologous prime-boost vaccination regimens may induce a balanced mucosal and systemic immune response in other vaccine platforms [49,50]. We therefore tested the ability of homologous IN/IN, homologous IM/IM, and heterologous IM/IN and IN/IM prime-boost vaccinations to stimulate both systemic and mucosal immune responses. Mice were primed either IM or IN with 5 µg H5 replicon complexed with NLC at an N:P of 10 and homologously or heterologously boosted 21 days post-prime (**Figure 6A**). As a negative control, animals were vaccinated IN/IN with 5 µg SEAP replicon complexed with NLC at an N:P of 10. At necropsy, 21 days post-boost, serum and BAL were collected to assay systemic and mucosal humoral responses, respectively, while spleens and lungs were collected to assay systemic and mucosal cellular responses, respectively.

**Figure 6.**
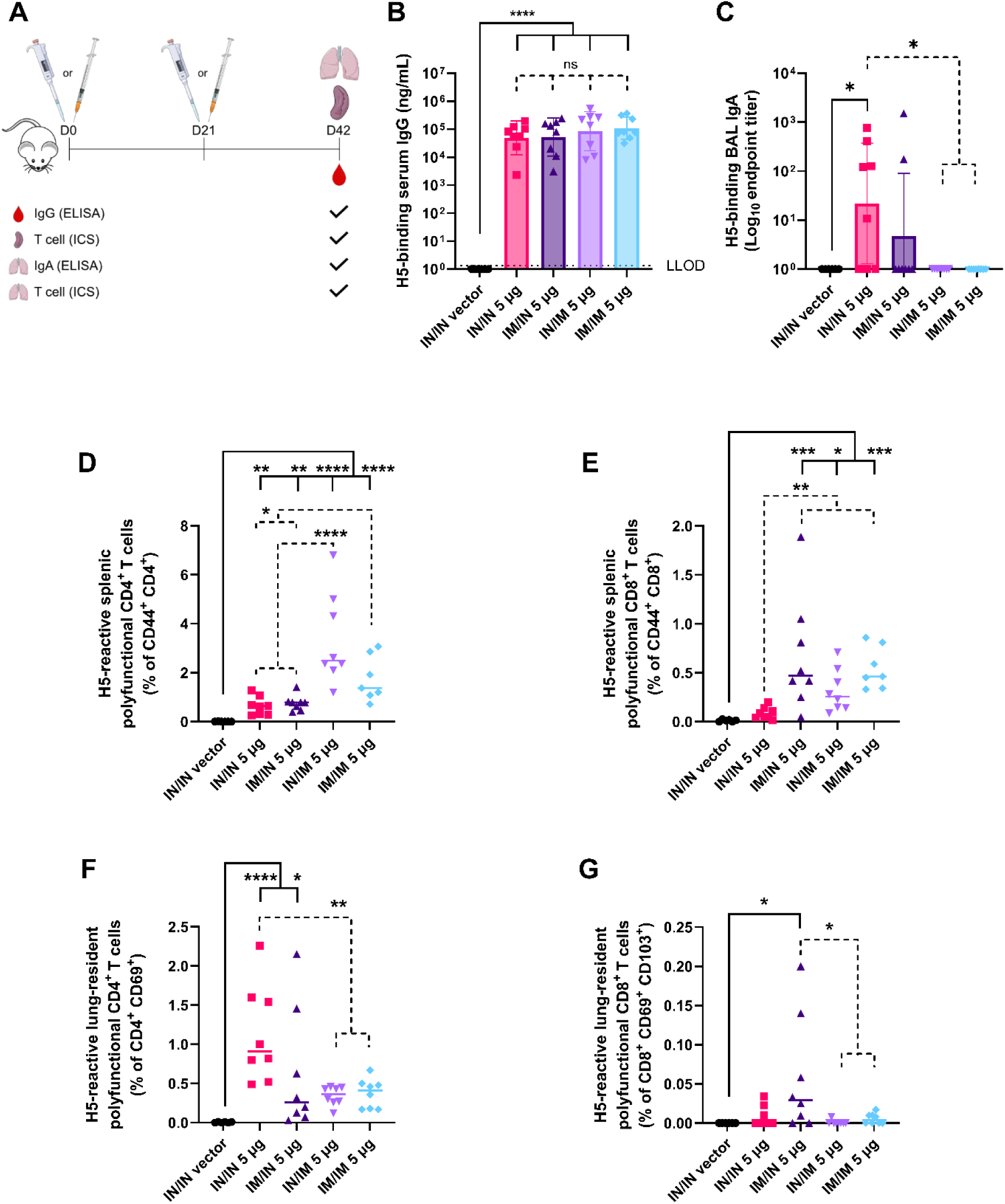
Heterologous dosing of H5 replicon vaccine candidate elicits a balanced immunological response. SEAP-expressing replicon was used as a negative vector control. All groups received a 5 µg dose of vaccine replicon. *n* = 8 mice (4 male and 4 female) per group except for SEAP control with *n* = 6 mice per group. (A) Heterologous dosing study design. (B-C) Antibody titers of mice after heterologous immunization with vaccine complex formed at N:P of 10. (B) H5-binding IgG post-boost serum titers. Data were log-transformed and displayed as the geometric mean ± geometric standard deviation. Data were analyzed using ordinary one-way ANOVA with Dunnett’s (solid line) or Tukey’s multiple comparisons test (dotted line). (C) H5-binding IgA post-boost bronchoalveolar lavage (BAL) titers. Data were analyzed using the Kruskal-Wallis test with Dunn’s multiple comparisons test (solid and dotted lines). (D-G) ICS analysis of polyfunctional T cells derived from mice after heterologous immunization with vaccine complex formed at N:P of 10. (D-E) H5-reactive polyfunctional (IFNγ^+^ IL-2^+^ TNFα^+^) splenic T cells. (F-G) H5-reactive polyfunctional (IFNγ^+^ IL-2^+^ TNFα^+^) lung-resident T cells. Horizontal lines show median. Data were log(y+1)-transformed and analyzed using one-way ANOVA with Dunnett’s (solid line) or Tukey’s (dotted line) multiple comparisons test. ns = not significant, * *p* < 0.05, ** *p* < 0.01, *** *p* < 0.001, **** *p* < 0.0001. IM = intramuscular, IN = intranasal, and LLOD = lower limit of detection.

Humoral immunogenicity was evaluated by ELISA for H5-binding serum IgG or BAL IgA titers. For serum IgG, all groups that received the H5 replicon produced a robust and similar H5-binding IgG response that was well above background, although mice vaccinated solely via the IM route showed a reduced spread in their response (**Figure 6B**). For BAL IgA, however, there was significant disparity between groups (**Figure 6C**). IN/IN vaccination elicited a strong though variable IgA response, with variability suspected to be due to variability of IN dosing in isoflurane-anesthetized mice. In contrast, the response in animals vaccinated IM/IM or IN/IM were indistinguishable from background (SEAP vector control). Interestingly, there was a BAL IgA response noted in some animals vaccinated IM/IN. Together, these data demonstrate that the H5 replicon vaccine is capable of generating robust systemic responses regardless of administration route and suggest that IN vaccination is required to induce mucosal immune responses.

Systemic and mucosal cellular immune responses were evaluated from spleens and lungs, respectively, and assayed by ICS and flow cytometry for H5-reactive polyfunctional T cells. Importantly, we observed that the systemic T cell response was strongly dependent on administration route, with splenic polyfunctional T cells observed most notably in animals vaccinated IM, either as a prime or boost dose (**Figure 6D-E**). In contrast, lung-derived polyfunctional CD4^+^ T cells were observed most notably in the IN/IN and IM/IN dosed groups, that is, those that received an IN boost dose of the H5 replicon vaccine (**Figure 6F**). Similar, though more subtle, results were observed for the lung-resident polyfunctional CD8^+^ T cell response (**Figure 6G**). Together, these data suggest that both IN and IM vaccine administration can stimulate systemic T cell responses, but a mucosal cellular response depends on at least one IN vaccine administration, preferentially when given as a boost dose.

### Intranasal prime-boost and intramuscular-prime, intranasal-boost replicon-NLC vaccination protect against viral-induced morbidity and mortality in ferrets

To assess the protective efficacy of the candidate vaccine formulations, we performed a lethal H5N1 viral challenge study in immunized fitch ferrets (**Figure 7A**). Ferrets were immunized with 10 µg H5 replicon-NLC prime-boost vaccinations 21 days apart, either homologously (IN/IN) or heterologously (IM/IN), or 10 µg SEAP replicon-NLC IN/IN as a vector control. Vaccinated animals were then intranasally challenged with a lethal dose of virus 21 days after receiving the boost vaccine dose. It was observed that both IN/IN and IM/IN H5 replicon-NLC vaccinated ferrets survived at an 83% rate, significantly surpassing the 17% survival rate of the vector control animals (**Figure 7B**). Clinical scores closely mirrored ferret survival (**Figure 7C**). While clinical scores of the vector-control vaccinated animals steadily climbed after viral challenge (ending in deaths of 5/6 ferrets), the two H5 replicon-NLC vaccine regimens were associated with significantly lower clinical scores with clear protection against virally-induced morbidity that exceeded that of the adjuvanted, inactivated whole virion control vaccine, despite weaker induction of serum antibody titers (**Supplementary** Figure 4). Compared to negative controls, viral loads in nasal washes from all vaccinated groups were significantly reduced 5 days post-infection, with near complete viral clearance in surviving vaccinated animals by study’s end (**Figure 7D**). Together, these data demonstrate the ability of our H5 replicon to provide robust infection-site protection against lethal H5N1 challenge following IN administration.

**Figure 7.**
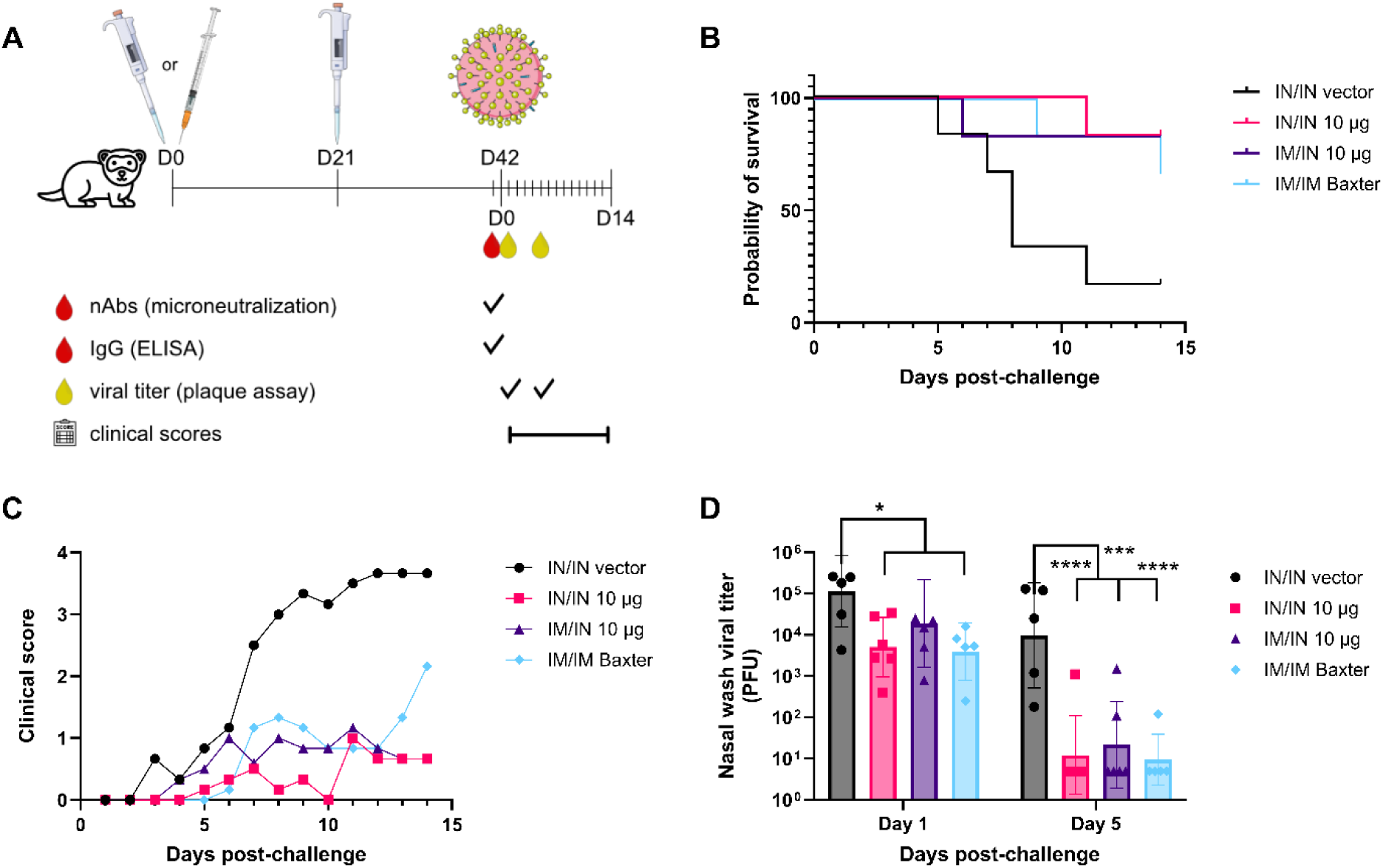
Efficacy of H5 replicon-NLC vaccine in ferrets. (A) Study design. Experimental animals were vaccinated with 10 µg H5 replicon either intranasally (IN) or intramuscularly (IM) and boosted intranasally 21 days later. *n* = 6 fitch ferrets (3 males and 3 females, 3 months of age) per group. Ferrets were challenged with 1 x 10^6^ PFU of A/Vietnam/1203/2004 H5N1 virus 21 days later. Baxter = positive-control alum-adjuvanted whole virion H5N1 vaccine. SEAP-expressing replicon was used as a negative vector control. Serum H5-binding IgG and neutralizing antibodies (nAbs) are shown in **Supplementary** Figure 4. (B) Survival plots of vaccinated ferrets after challenge. (C) Average clinical score of vaccinated ferrets after challenge. (D) Viral load in nasal wash post-challenge (geometric mean and geometric SD). Data were log-transformed and analyzed using a mixed-effects analysis with Dunnett’s multiple comparisons test. * *p* < 0.05, *** *p* < 0.001, **** *p* < 0.0001.

## DISCUSSION

Intranasal vaccines have the important attribute of being able to induce a mucosal immune response at the primary site of infection, which has the benefit of limiting viral uptake, replication, and transmission of respiratory viruses, as well as helping to alleviate vaccine hesitancy through needle-free delivery [19,31,51]. For many vaccines, conversion from IM to IN administration requires extensive vaccine reformulation in order to achieve sufficient mucoadhesion and mucopenetration for efficient uptake within the nasal cavity [52]. In the present work, we utilized an NLC platform to develop an H5 influenza replicon vaccine candidate that can be administered IN or IM and achieve a high level of protection from viral-associated disease without the need for vaccine reformulation between the two administration routes.

NLC-based vaccine complexes are structurally distinct from LNP-based vaccines. More specifically, a nucleic acid is encapsulated within the core of LNPs [53], while here the replicon is electrostatically adsorbed to the surface of NLCs [26], in a manner similar to the formation of cationic nanoemulsion (CNE)-based vaccines [54]. As such, known structural insights for LNP and CNE-based nucleic acid vaccines are not necessarily applicable to replicon-NLC vaccine structures, and in-depth characterization of the replicon-NLC complex is necessary to develop an appropriate IN vaccine. In particular, the relationship between the N:P ratio and the biophysical characteristics of the H5 replicon-NLC vaccine must be understood, as this ratio affects the apparent zeta potential, size, concentration, and degree of replicon protection for the vaccine, thus affecting its efficacy. Here, we observed that an N:P ratio ≥ 5 was necessary to achieve a high degree of protection of the H5 replicon construct from enzymatic degradation, and that this corresponded with a positive bulk zeta potential. At N:P ratios ≤ 1, the measured bulk zeta potential of the complex became negative, and replicon protection was concomitantly lost, suggesting increased bulk solvent exposure of the vaccine replicon at low N:P ratios due to steric interference at the NLC surface from multiple replicon molecules, as indicated by TEM imaging. This indicates that the N:P ratio directly dictates the structure of the final replicon-NLC vaccine complex (**Figure 8**): at high N:P ratios, there is a relative excess of NLCs that enables complete binding of the entirety of the replicon nucleic acid by multiple NLC particles, inducing the formation of a pseudo- capsule-like structure around the replicon, which sequesters it from the bulk solution and confers protection from enzymatic degradation. However, at low N:P ratios, there becomes a relative excess of replicon nucleic acid such that the NLCs present in solution are unable to bind and sequester the entire length of the replicon sequence, leaving the replicon exposed to the bulk solution and accessible for enzymatic degradation. The measured zeta potentials, cationic at high N:P ratios and anionic at low N:P ratios, reflect this organization.

**Figure 8.**
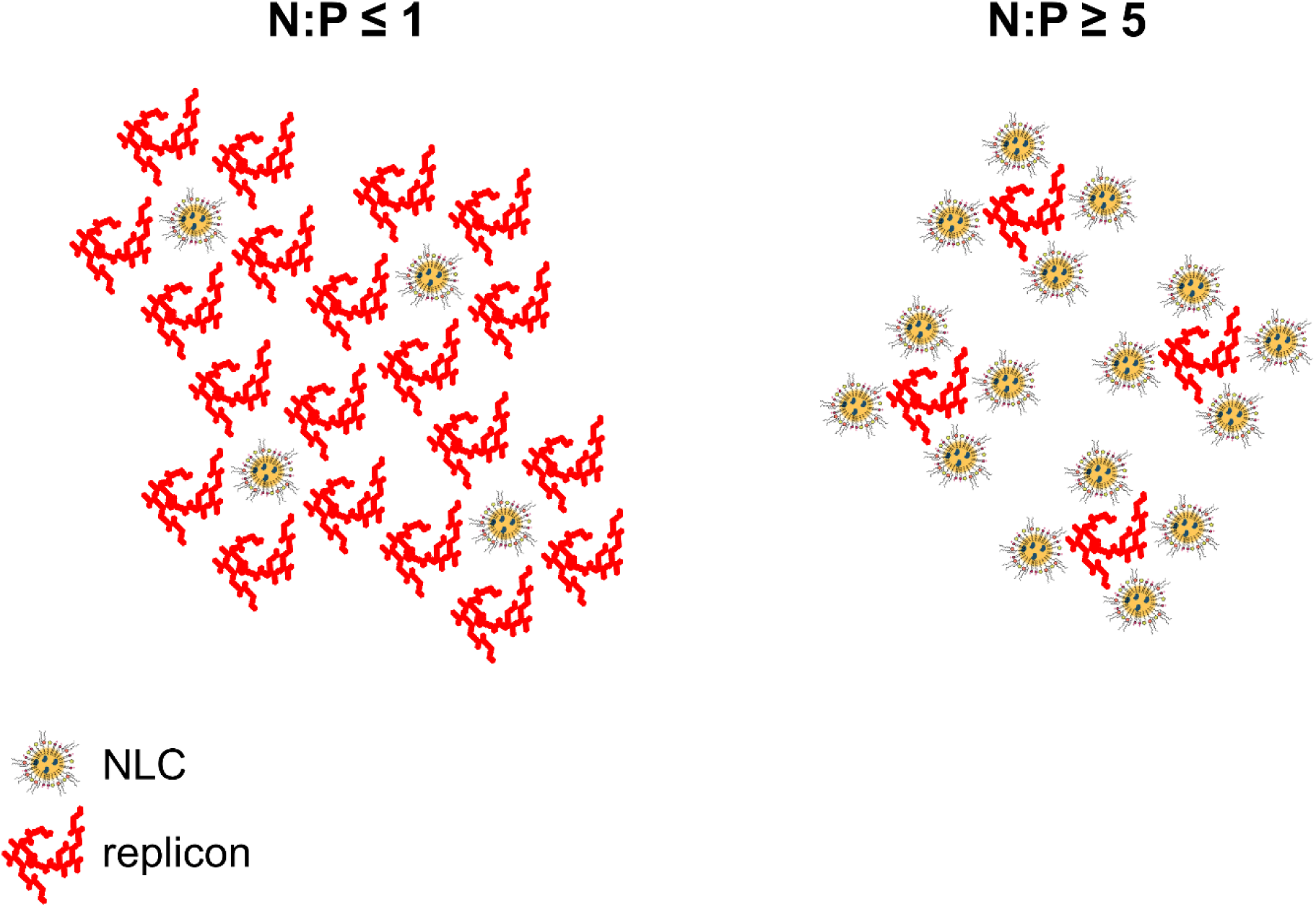
Model of replicon-NLC complexes at different N:P ratios. Simple mixing of NLC with replicon produces an ensemble of vaccine complexes. As the N:P ratio increases to ≥ 5, the number of well-formed/functional complexes increases, resulting in an increase in gene-delivery events and downstream immunogenicity. Formulations with low N:P ratios ≤ 1 have markedly fewer functional complexes and thus do not induce a sufficient immune response in mice. Well-formed complexes are characterized by a positive zeta potential and ability to protect RNA from enzymatic degradation.

The apparent hydrodynamic size of the complexes was also influenced by the N:P ratio, with higher N:P ratios generally resulting in a smaller overall Z-average particle size as measured by DLS. The DLS size distributions appeared to reflect intrinsic heterogeneity upon complexing of the NLCs with the replicon, in contrast to LNP-based complexes, which tend to be more homogeneous in nature. This heterogeneity was confirmed using NTA and cryo-TEM. The advantage of this heterogeneity unique to the NLC platform is that it allows for more complete pseudo-encapsulation and protection of the replicon. Others have previously shown that it is extremely difficult to encapsulate lengthy nucleic acids within LNPs while maintaining a high degree of replicon sequence integrity [53]. In contrast, because multiple NLCs can surround an individual replicon molecule, at an appropriate N:P ratio, it is possible to achieve more complete protection of the replicon to allow for a higher effective dose of replicon to be delivered upon IN or IM administration.

At all tested N:P ratios, the nanoparticle size distribution spanned from approximately 40 to 250 nm. Total nanoparticle concentration decreased as the N:P ratio was decreased down to an N:P of 1. It was notable that the overall trend in nanoparticle concentration appeared to correlate well with the degree of apparent vaccine uptake in Vero cells. Considering the structures observed by TEM indicating extensive replicon exposure to the bulk solvent at a low N:P ratio, and given that it is typically difficult for a nucleic acid alone to cross the plasma membrane without the engagement of active cellular transport mechanisms [55], this result suggests there is an active (i.e., functional) vaccine complex that forms in the 40 – 250 nm size range. These cationic functional vaccine complexes are thus hypothesized to deliver replicon to cells by endosomal destabilization via the proton-sponge effect and release of the replicon cargo into the cytosol, in a mechanism consistent with that of other cationic lipid platforms used for nucleic acid delivery [46]. We note that some vaccine uptake was also observed at N:P ratios of 1 and 0.6; it is likely that even at these low N:P ratios, due to the heterogeneity of the system, there is a small population of replicon-NLC complexes not captured in the measured zeta potential of the bulk formulation that forms with a positive surface charge and is able to enter and deliver nucleic acid to cells. However, *in vivo* studies demonstrate that this small fraction of functional complex at N:P ratios ≤ 1 is insufficient to stimulate immune responses. Vaccine IgG and CD4^+^ and CD8^+^ T cell responses largely reached a plateau at N:P ratios of 8 – 15, with an intermediate level of immunogenicity observed at an N:P of 5. Taken together, the biophysical and immunological data suggest that we can elicit a sufficient immune response when dosing IN within the N:P range of 8 – 15 and suggest that vaccine nanoparticle concentration plays a crucial role in vaccine efficacy. We note that N:P ratios above 15 were not explored in the present work due to previous studies suggesting no additional immunological benefit at higher N:P ratios [26]. Subsequent *in vivo* dosing and challenge studies with the H5 replicon-NLC vaccine complexes were carried out using an N:P of 10 to ensure we remained above the minimum N:P ratio needed to achieve high immunological responses while simultaneously minimizing the net DOTAP content and thus risks of potential DOTAP-related IN vaccine reactogenicity.

To evaluate the efficacy of the N:P ratio-optimized IN vaccine, various homologous and heterologous prime-boost dosing strategies were evaluated in mice for their ability to induce adaptive and innate mucosal immune responses. It is known that IgA antibody and CD4^+^ and CD8^+^ T cell-mediated mucosal immune responses play important roles in the prevention of early-stage viral infection replication and disease transmission [19,56–58]. In the present work, we observed that IN/IN and IM/IN dosing of the H5 replicon-NLC vaccine complex at a 5 µg replicon dose uniquely induced mucosal humoral immune responses in addition to systemic responses. Indeed, the induction of both strong systemic and mucosal immune responses by the IM/IN dosing strategy suggests a potential benefit of using a heterologous IM/IN dosing strategy, consistent with previous observations when using an IM/IN dosing strategy [8,35].

Having established a mucosal immune response using IN/IN or IM/IN dosing strategies, we next sought to investigate the protective efficacy of the H5 replicon-NLC vaccine using a ferret challenge model of H5 influenza viral infection. Using a moderate replicon dose (10 µg), the IN/IN or IM/IN administered vaccine protected animals from morbidity and mortality and led to rapid control of viral loads, superior to the performance of an IM/IM Baxter vaccine control and previously published IM replicon influenza vaccines [59]. This excellent vaccine performance may be due to the inclusion of IN vaccination with a replicating nucleic acid that mimics a viral infection and stimulates robust innate immunity at mucosal sites [60,61], with resulting mucosal immune response at the primary site of infection as demonstrated in the mouse model but unmeasured in ferrets. Taken together, these data demonstrate that the IN administered H5 replicon-NLC vaccine affords a high level of protection while completely obviating the need for needle-based delivery.

There are a few limitations of this study that should be noted. First, only one antigen was tested, the full-length HA protein, with no alternatives or comparators. Despite this, others have demonstrated the full-length, membrane-anchored HA functioned better than various other secreted, truncated, or multimerized forms of HA [62], rendering this a suitable replicon system for these studies. It should also be noted that IN dosing in smaller animals (e.g., mice, ferrets, etc.) can be somewhat variable and likely contribute to some of the observed spread in immune response data in IN dosed animals. Ultimately, vaccine efficacy when administered IN will need to be assessed in larger animals with similar anatomical physiology to humans, such as non-human primates. Further, due to funding limitations, the work performed here did not include a full dose-response assessment of the replicon vaccine in the ferret model to identify the optimal IN dose. Increasing the nominal IN dose has since been demonstrated to further enhance protection in ferrets to complete protection against morbidity and mortality [35]. Finally, we note that we were unable to provide a direct comparison against an optimal H5 vaccine control. The positive control available and used here, an IM administered Baxter alum-adjuvanted whole virion H5N1 vaccine, does not reflect the current standard of care for influenza vaccines containing adjuvants such as AS03 or MF59, and comparison of the IN administered replicon vaccine with another IN vaccine was not possible due to the lack of any such H5 vaccine developed to date suitable for IN administration.

## CONCLUSIONS

We have successfully developed a replicon vaccine capable of protecting against lethal H5N1 influenza viral infection and inducing robust systemic and mucosal immune responses when delivered by IN administration in mice and ferrets. Moreover, we have shown that the same vaccine formulation can be delivered both IM and IN, with mucosal immune responses uniquely induced by IN vaccination. We further found a mixed IM-prime, IN-boost dosing strategy may be the most beneficial for eliciting the strongest balanced systemic and mucosal immune responses, and the broadest protective responses in individuals. This has important implications for the development of effective influenza and other respiratory viral vaccines as the existence of a unified formulation for both systemic and mucosal dosing provides unparalleled flexibility to allow for bespoke dosing strategies tuned to the disease in question. Furthermore, by eliminating the need for route-dependent vaccine reformulation, we substantially decrease the cost (both monetary and time) of manufacturing and distributing vaccines in the event of the next viral outbreak.

## ACKNOWLEDGEMENTS

Cryo-TEM images were collected by NanoImaging Services (San Diego, CA, USA). Images for figures were obtained from Clker.com under a CC0 1.0 license (https://creativecommons.org/publicdomain/zero/1.0/); amoghdesign and Mihimihi at Freepik (https://www.freepik.com/); Nicolás De Francesco at SciDraw (https://scidraw.io/) under a CC BY 4.0 license (https://creativecommons.org/licenses/by/4.0/); Ryan Kissinger at NIH BIOART Source (https://bioart.niaid.nih.gov/bioart/150, https://bioart.niaid.nih.gov/bioart/187, and https://bioart.niaid.nih.gov/bioart/279); and Servier Medical Art (https://smart.servier.com/) under a CC BY 4.0 license (https://creativecommons.org/licenses/by/4.0/). The following reagent was obtained through BEI Resources, NIAID, NIH: A/H5N1 Influenza Vaccine, Inactivated Whole Virion (A/Vietnam/1203/2004), Vero-Cell Derived, Adjuvanted, 15 Micrograms HA, NR-12143. We thank Jeffrey A. Guderian for their assistance with the manufacturing of the replicon constructs used for these studies, and we thank Dr. Michael Davis and Dr. Valerie Soza for editing and proofreading the manuscript.

## AUTHOR CREDIT STATEMENT

**Wynton D. McClary:** conceptualization, formal analysis, investigation, methodology, writing – original draft, writing – review & editing. **Devin S. Brandt:** conceptualization, data curation, formal analysis, investigation, methodology, project administration, software, supervision, validation, visualization, writing – original draft, writing – review & editing. **Madeleine F. Jennewein:** conceptualization, investigation, data curation, methodology, project administration, supervision, visualization, writing – review & editing. **Jasneet Singh:** investigation, writing – review & editing. **Samuel Beaver:** investigation, writing – review & editing. **Matthew R. Ykema:** investigation, writing – review & editing. **Christopher Press:** resources, writing – review & editing. **Eduard Melief:** resources, writing – review & editing. **Julie Bakken:** investigation, writing – review & editing. **Pauline Fusco:** investigation, writing – review & editing. **Ethan Lo:** investigation, writing – review & editing. **Peter Battisti:** investigation, writing – review & editing.

**Noah Cross:** investigation, writing - review & editing. **Darshan N. Kasal:** data curation, writing – review & editing. **Airn Tolnay Hartwig:** investigation, writing – review & editing. **Corey Casper:** funding acquisition, writing – review & editing. **Richard A. Bowen:** conceptualization, funding acquisition, project administration, resources, supervision, writing – review & editing. **Alana Gerhardt:** conceptualization, funding acquisition, supervision, project administration, methodology, writing – review & editing. **Emily A. Voigt:** conceptualization, funding acquisition, methodology, project administration, supervision, writing – review & editing.

## DATA AVAILABILITY

Data will be made available on request.

