## Supplementary Material for "Development of an Intranasally- and Intramuscularly-Administrable Replicon Vaccine Efficacious Against H5N1 Influenza Virus"

**Supplementary Table 1.** Zeta potential of H5 replicon-NLC complexes

| N:P Ratio | Zeta Potential (mV) |
|---|---|
| 15 | 14.0 ± 0.5 |
| 12 | 13.5 ± 0.4 |
| 10 | 13.3 ± 0.5 |
| 8 | 13.7 ± 0.2 |
| 5 | 14.6 ± 0.1 |
| 1 | -36.9 ± 0.5 |
| 0.6 | -53.9 ± 1.9 |

**Supplementary Table 2.** Z-average diameter and polydispersity index (PDI) of H5 replicon-NLC complexes

| N:P Ratio | Z-Average Diameter (nm) | PDI |
|---|---|---|
| 15 | 116.9 ± 1.1 | 0.508 ± 0.005 |
| 12 | 135.9 ± 3.3 | 0.530 ± 0.019 |
| 10 | 137.4 ± 3.9 | 0.490 ± 0.018 |
| 8 | 144.6 ± 3.5 | 0.484 ± 0.012 |
| 5 | 193.4 ± 6.8 | 0.449 ± 0.042 |
| 1 | 1395.0 ± 195.2 | 0.736 ± 0.041 |
| 0.6 | 452.4 ± 13.2 | 0.472 ± 0.036 |

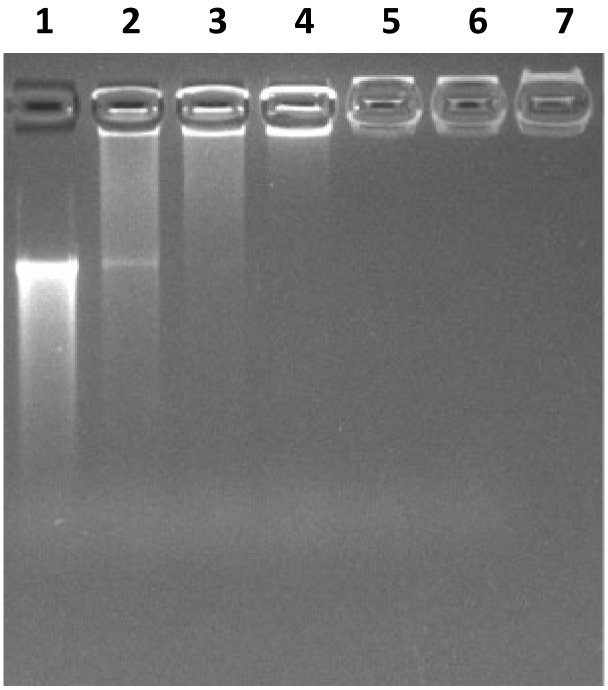


**Lanes:**

1. **Free replicon**
2. **N:P 0.6 – Whole Complex**
3. **N:P 0.8 – Whole Complex**
4. **N:P 1 – Whole Complex**
5. **N:P 3 – Whole Complex**
6. **N:P 5 – Whole Complex**
7. **N:P 15 – Whole Complex**

**Supplementary Figure 1.** Maintenance of H5 replicon nucleic acid binding to NLCs. Replicon-NLC vaccine complexes were loaded directly onto a 1% agarose gel without organic extraction of the nucleic acid. Replicon that remains in complex with the NLC will not migrate into the agarose gel lanes due to the maintained interaction with the lipids and is indicative of the maintenance of the complete replicon-NLC vaccine complex. Vaccine complexes were assessed from a N:P range of 0.6 (saturating replicon) to 15.

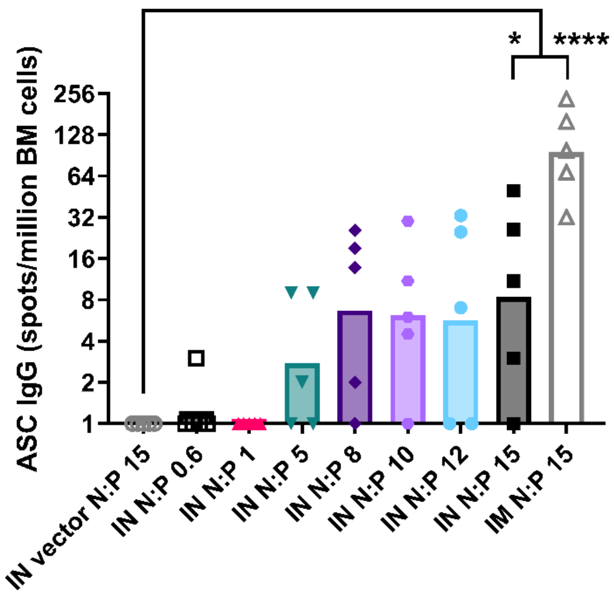


**Supplementary Figure 2.** Bone marrow IgG ELISpot analysis of antibody-secreting cells (ASC) harvested from mice that were dosed with H5 replicon vaccine complexes formed at various N:P ratios. Related to Figure 5. A complex formed at N:P of 15 using a SEAP-expressing replicon was used as a negative vector control. $n = 5$ mice per group (C57BL/6J female mice, 6 to 8 weeks of age). Data were log-transformed and analyzed using one-way ANOVA with Dunnett's multiple comparisons post-hoc test. * $p < 0.05$, **** $p < 0.0001$. Data displayed as geometric mean. BM = bone marrow, IM = intramuscular, IN = intranasal.

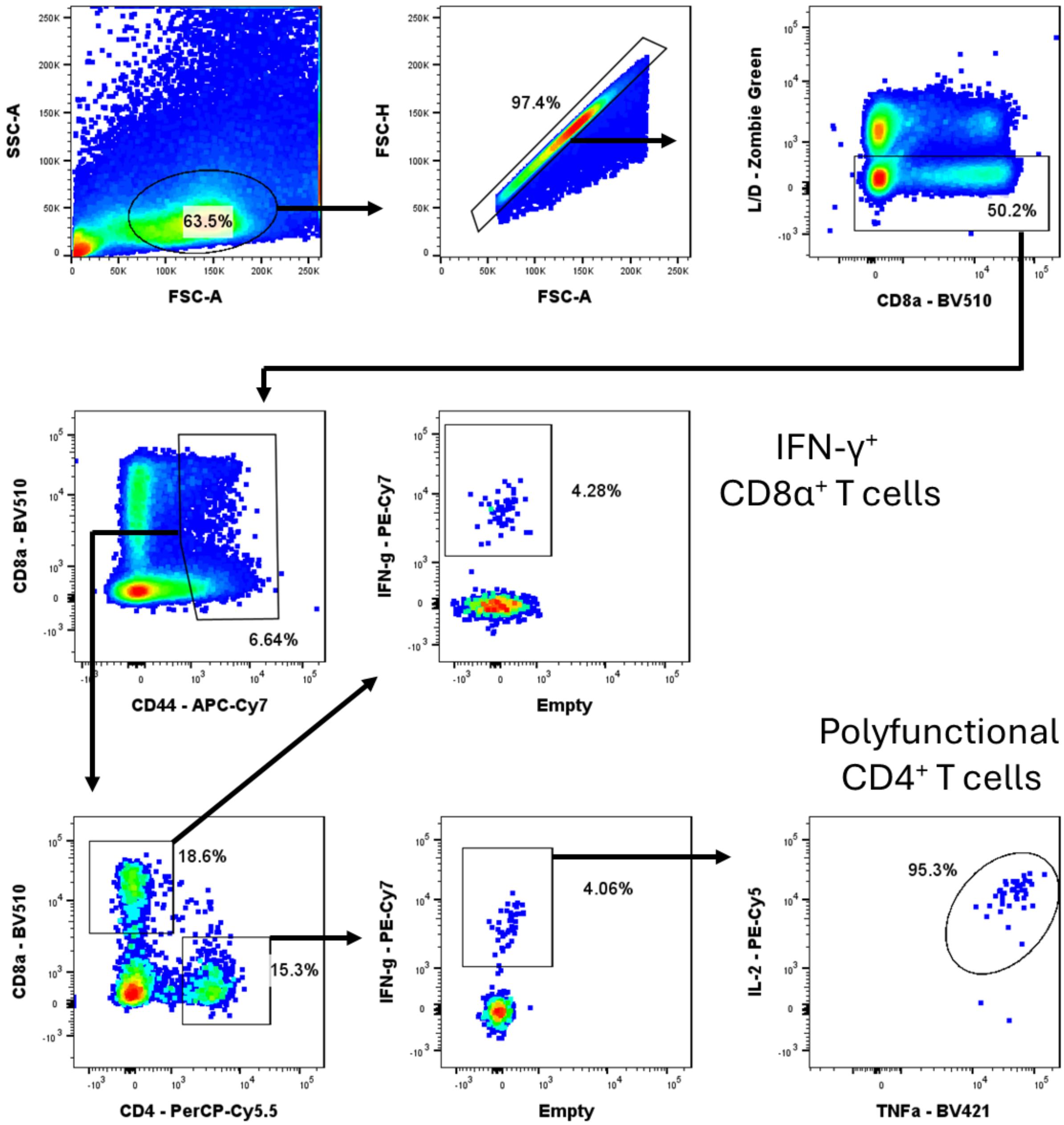


**Supplementary Figure 3.** Flow cytometry gating strategy for splenic antigen-responsive T cells. Spleens were processed and labelled as indicated in the Materials and Methods section. Gating outlines the stepwise selection process used to identify IFN-$\gamma^+$ CD8$\alpha^+$ T cells and polyfunctional CD4$^+$ T cells.

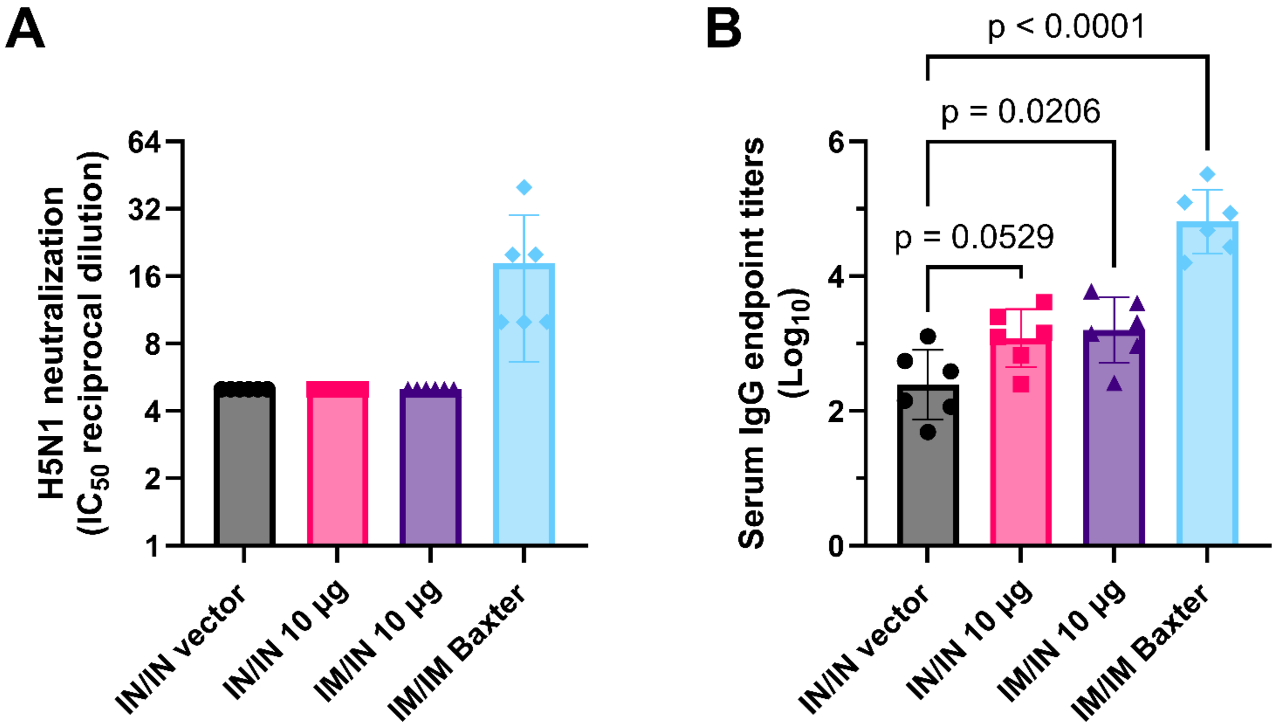


**Supplementary Figure 4.** Neutralizing and H5-binding antibody titers in vaccinated ferret sera. Related to Figure 7. Experimental animals were vaccinated with 10 µg H5 replicon-NLC either intranasally (IN) or intramuscularly (IM) and boosted intranasally 21 days later. Serum was collected just prior to (Day -1) challenge. $n$ = 6 fitch ferrets (3 males and 3 females, 3 months of age) per group. Baxter = positive-control alum-adjuvanted whole virion H5N1 vaccine. SEAP-expressing replicon was used as a negative vector control. (A) Serum neutralizing antibody titers as measured via influenza microneutralization assay. (B) Serum H5-binding IgG endpoint titers. Data were analyzed using ordinary one-way ANOVA with Dunnett's multiple comparisons test. Data presented as the mean ± standard deviation.
